# VSTseed: periodic spaced seeds for reads with substitutions

**DOI:** 10.1101/2021.06.09.447791

**Authors:** Valeriy Titarenko, Sofya Titarenko

**Affiliations:** School of Biological Sciences, University of Manchester, Manchester, M13 9PL, UK; School of Computing and Engineering, University of Huddersfield, Huddersfield, HD1 3DH, UK

## Abstract

**Motivation:** Technical progress in computer hardware made it possible to access and process large amounts of data even on budget workstations. Therefore new or existing alignment algorithms may use large index files to increase performance. Spaced seeds with large weights reduce the number of possible locations of a read within a reference sequence. Optimal patterns for spaced seeds may guarantee to align reads even with several substitutions.

**Results:** For reads of 64–200 bp periodic spaced seeds of 32, 40, 48, 56, 64 weights are found that guarantee to locate all positions within a reference sequence for a specified number of point mutations. SIMD instructions to convert masked reads into 64, 80, 96, 112, 128-bit numbers are provided.

**Availability:** C codes to generate spaced seeds and find optimal SIMD instructions for them are freely available under MIT license at https://github.com/vtman/VSTseed

## 1 Introduction

The most common question in sequence analysis is whether two sequences are related. To answer this question we need to align the sequences or their parts. For the past two decades several sequence mapping and alignment software packages based on statistical methods were released and earned their popularity, e.g. BLAST [1], Bowtie2 [15], BWA-MEM/SW [18,19], SHRiMP2 [9], MiniMap2 [17]. While these algorithms or their modifications are still effective and very popular, they were created when significant restrictions on computational resources were in place. Initially amounts of data available for scientists for each experiment were relatively small, so computers were able to process them. However, with better experimental hardware for sequencing sizes of datasets started to increase very fast. As the result some approaches for sequence alignment could not be used for real problems with large input datasets. These days we may observe significant changes in the size of random-access memory and read/write speeds for storage devices available even for a budget computer. Therefore many issues related to the size of memory and speed of access to storage data became obsolete. As the result modern algorithms and software tools need to overcome different issues to improve performance, e.g. organise proper data alignment to exploit SIMD (Single Instruction, Multiple Data) instructions and fast access to CPU’s cache, parallelise processing for multithreading applications.

The most recent human reference genome assembly (GRCh38.p13, released in 2019) contains *L* = 3,272,116,950 ≈ 3.3 · 10^9^ bases in total. This means that each position within the reference sequence can be written as a 32-bit unsigned integer (2^32^ ≈ 4.3 · 10^9^ > *L*). For the human reference genome each base either belongs to the alphabet ∑ = {**A, C, G, T**} or is symbol **N** which is any of the four bases. Suppose there is a sequence of *K* integer numbers of positions (all different). Then for each position in the reference genome we get *K* bases according to the given integer sequence. If there are no symbols N among the selected *K* bases, then this *K*-sequence of bases is identified by 2*K*-bit “signature” (2 bits for each base). So, for each position in the reference sequence we may store 32 bits for its position and 2*K* bits for the “signature”, or *L*(32 + 2*K*) bits in total for the whole reference sequence. The list of pairs (“position”, “signature”) can be sorted by “signature” values and be grouped in several arrays. For example, based on values of first 16 bits of “signatures” we may split the list into 65536 = 2^16^ arrays and omit the first 16-bits for each pair as they are all the same for pairs in the arrays. Thus to store the list we may need only around *M* = *L*(16 + 2*K*) bits. If *K* = 32, then *M* ≈ 3.3 · 10^9^ · (16 + 64)/8 ≈ 33.0 GB, and for *K* = 64, we get *M* ≈ 59.4 GB. Once pre-calculated the list of pairs can be stored on a hard drive and accessed within several seconds by a sequence alignment algorithm.

After data collection a researcher is provided with a list of a fixed-length sequences of bases (*reads*). A sequence alignment algorithm tries to find best possible positions within the reference sequence such that a distance between a read and a substring of the reference sequence is minimal. The distance between two strings *x* and *y* was introduced in [11] and can be determined in several ways. We may define a list of elementary operations which convert a source string *x* into a target string *y*. Each operation may have a different cost, the distance is then a sum of all costs. If we cannot transform *x* into *y*, then the cost is ∞. According to [23], depending on permissible elementary operations the most common distances are: Levenshtein (or edit) distance (insertions, deletions, substitutions are allowed at equal cost of 1; symmetric, i.e. *d*(*x, y*) = *d*(*y, x*)) [16], Hamming distance (only substitutions, symmetric) [13], episode distance (only insertions, is not symmetric) [8], longest common subsequence distance (insertions and deletions, all costing 1, symmetric) [3, 24].

Similarity between two strings were originally measured using dynamic programming algorithms, see [10]. However, due to time complexity the use of these algorithms became impractical for the increased size of data available. In late 90s several new algorithms, e.g. BLAST [1, 2], appeared based on ideas of filtration and indexing. Short sequence fragments (*seeds*) were used. To align a read we require seeds from the read to be also found in a reference sequence. The search is sped up using various indexing, e.g. hash-tables, which may provide a researcher with “false-matching” positions. The seeds were considered as contiguous segments of 1s (1 is when corresponding bases for the search and target sequences are compared, and 0 when they are ignored). In PatternHunter [21] spaced seeds were used, e.g. for weight 11 (total number of 1s) the most sensitive seed was `111010010100110111`. Some spaced seeds for different weights can be found in [6]. The idea of spaced seeds were extended for other problems: vector [4], indel [22], neighbor [7] seeds. In ZOOM software [20] spaced seeds are generated to perform alignment with at most 2 mismatches. In PerM software [5] so-called periodic spaced seeds are used to improve efficiency of mapping.

In this paper we also consider optimal spaced seeds as periodic ones, however unlike similar problems solved in other software, e.g. PerM [5], RMAP [26], ZOOM [20], SeqMap [12], we focus on seeds that are designed for reads of 100–250 bp, have weights between 32 and 64, can tolerate up to 6 single letter mismatches. Seeds with large weights allow to decrease the number of candidate positions within a reference sequence, thus helping to improve performance of an alignment algorithm. We also provide software tools to generate the seeds for reads of a given length and a known maximum number of substitutions, convert the spaced seeds into contiguous arrays (in order to generate “signatures”) using SIMD instructions. Once a spaced seed has been found, the software can provide a user with index files (30–60 GB in total) within 2–3 minutes on a budget computer.

## 2 Materials and methods

### 2.1 Digital representation of sequences

Suppose we have a reference sequence of length *L* and a set of reads of length *n*. For the human reference genome each base either belongs to the alphabet ∑ = {**A, C, G, T**} or is symbol **N**. To perform sequence alignment the original strings should be stored in a computer’s hard drive/memory. Each of 5 symbols may be encoded with numbers 0, …, 4. We may combine *k* consequent symbols and encode them *m*-bit numbers such as 5^*k*^ ≤ 2^*m*^. If *k* = 1, then *m* = 3 and numbers 5, 6 and 7 are not used, so this storage scheme requires extra 3/5 = 60% of space compared to the ideal case. If *k* = 2, then *m* = 5 and extra space is (32 − 25)/25 = 28%; for *k* = 3 we get *m* = 7 and 2.4% (the best value for *k* < 27). So, if no additional lossless compression is preformed, then coding each 3 consequent bases with 7-bit numbers is close to the ideal case (Supplementary Material S1).

When we have two sequences of the same length and want to find the corresponding Hamming distance (or, similarly, count the number of same bases at same positions), we may, of course, perform comparison per each base, however to improve performance the use of 128-, 256-, 512-bit SIMD instructions can be beneficial. If at same positions of two sequences there are letters of the alphabet ∑, then the Hamming distance does not change if the letters are the same, or increased by one, otherwise. However, dealing with symbol **N** may be more complicated. As symbol **N** may have the same chance to be one of the symbols of ∑, then we may set the Hamming distance to *d*(*S*, **N**) = 3/4 for any symbol *S* ∈ ∑. At the same time we may also set *d*(*S*, **N**) = 1. Similarly, there may be three options for *d*(**N, N**): 0, 1 or 3/4. So, processing symbol **N** differently from the symbols of the alphabet ∑ may further decrease performance.

We propose the following digital representation of a sequence. Each 32 consequent symbols are represented as 128-bit numbers, i.e. one symbol requires 4 bits. Thus the extra space is (16 − 5)/5 = 220%, so the storage scheme is very inefficient. On the other hand, storing data for the human reference genome even in uncompressed format requires only 1.65 GB, i.e. a fraction of RAM available in budget computers these days. Therefore the storage overhead should not be a bottleneck for a modern computer. However, the proposed scheme may provide us with performance benefits.

Let us agree to number all positions from 0, i.e. the first symbol has index 0. A bit from the first 32-bit chunk (bits [0, 31]) is set to 1 if the corresponding symbol of the sequence is A, otherwise it is set to zero. In a similar way we set bits [32, 63] (symbol C), bits [64, 95] (symbol G) and bits [96, 127] (symbol T). According to this definition only one of four bits *i*, (*i* + 32), (*i* + 64) and (*i* + 96) can be 1. By settings all four bits to zeros we define symbol **N**. If we have two sequences of 32 symbols we may write them as `__m128i` structures *a* and *b* used for Intel Intrinsic instructions (Supplementary Material S2). We just need one bitwise AND instruction (`_mm_and_si128`) to check if corresponding bits has 1s at the same positions and then count the total number of 1s with shift, bitwise OR and `popcnt` instructions.

~~~
**int inum, ires;
__mm128i a, b, u1, u2, u3, u4, u5;
u1 = _mm_and_si128(a, b);
u2 = _mm_bsrli_si128(u1, 8);
u3 = _mm_or_si128(u1, u2);
u4 = _mm_bsrli_si128(u3, 4);
u5 = _mm_or_si128(u3, u4);
inum = _mm_extract_epi32(u5, 0);
ires = _mm_popcnt_u32(inum);**
~~~

As the result we get a number of positions with the same symbols of the alphabet ∑ (Supplementary Material S3). The Hamming distance is then `(32-ires)` under assumption of *d*(*S, S*) = 0, *d*(*S*, **N**) = 1 and *d*(**N, N**) = 1.

### 2.2 Indexing of the reference genome

The above 128-bit representation of 32-symbol sequences can also be used for indexing of the reference genome by splitting it into chunks of 32 symbols (with possible padding by symbols **N**). Thus we get a sequence of 128-bit numbers. Using bitwise shift and logical instructions it is possible to find the 128-bit number corresponding to any starting position (Supplementary Material S3) of the reference genome. Comparing `inum` with `0xFFFFFFFF` we may avoid chunks containing symbols **N**. In this case the 128-bit number can be converted to a 64-bit number. For this purpose we may use the logical Table 1 and the following SIMD instructions

**Table 1.** Truth table for A, C, G, T, A|C, and C|G.

| A | C | G | T | A\|C | C\|G |
| --- | --- | --- | --- | --- | --- |
| 1 | 0 | 0 | 0 | 1 | 0 |
| 0 | 1 | 0 | 0 | 1 | 1 |
| 0 | 0 | 1 | 0 | 0 | 1 |
| 0 | 0 | 0 | 1 | 0 | 0 |

~~~
**__mm128i mACGT, mCGT0, mres;
mCGT0 = _mm_bsrli_si128(u1, 4);
mres = _mm_or_si128(mACGT, mCGT0);**
~~~

Then the first 32 bits of `mres` corresponds to A|C values (symbol “|” means bitwise OR) and the second 32 bits are for C|G, so the first 64 bits can be used to create an index file for the reference genome.

Each position within a human reference genome can be coded as a 32-bit integer number. Therefore a list of pairs (“position”, 32-sequence) requires 32 + 64 = 96 bits for each pair, since we have only 4 symbols and require 2 bits for each symbol to form 64-bit “signature” number. The list contains around 3.3 · 10^9^ pairs and can be sorted by “signature” values. For a given read we may find all its 32-symbol subsequences and by using the list we obtain all positions within the reference genome where the read and the reference genome have at least one common contiguous 32-symbol chunk. We may reduce the size of memory required to store the list by splitting the list into 65536 = 2^16^ arrays, so instead of 64 bits for “signatures” we may store only 64 − 16 = 48 bits or 10 bytes per pair. In this case we need about 3.3 · 10^9^ × 10 = 33 GB. If a similar approach is applied to 64-symbol chunks, then the size of memory becomes 59.4 GB. Many modern motherboards have memory capacity of at least 128 GB. Index files take some time to generate, however they need to be created only once for a given reference genome and a chosen length of a substring. Accessing these files is straightforward and takes several seconds with modern storage drives, e.g. budget NMVe (Non-Volatile Memory Express) devices have read speeds of 5 GB/s.

However, there are two main bottlenecks. Firstly, for some subsequences there may be too many candidate positions. We have applied the above procedure for the human genome and used contiguous chunks of various lengths. The percentage of unique “signatures” varies from 78.5% for 32-symbol chunks to 86.7% for 64-symbol chunks. However, there are “signatures” occurring more than thousands times. For a given integer number we may find percentage of all “signatures” that have more occurrences than the given number (see Figure 1). So, if we randomly pick up a contiguous 32-symbol sequence, then there is 5% chance that it has more than 200 positions within the reference genome. For 64-symbol sequence, the chance is reduced to 1.67%.

**Figure 1.**
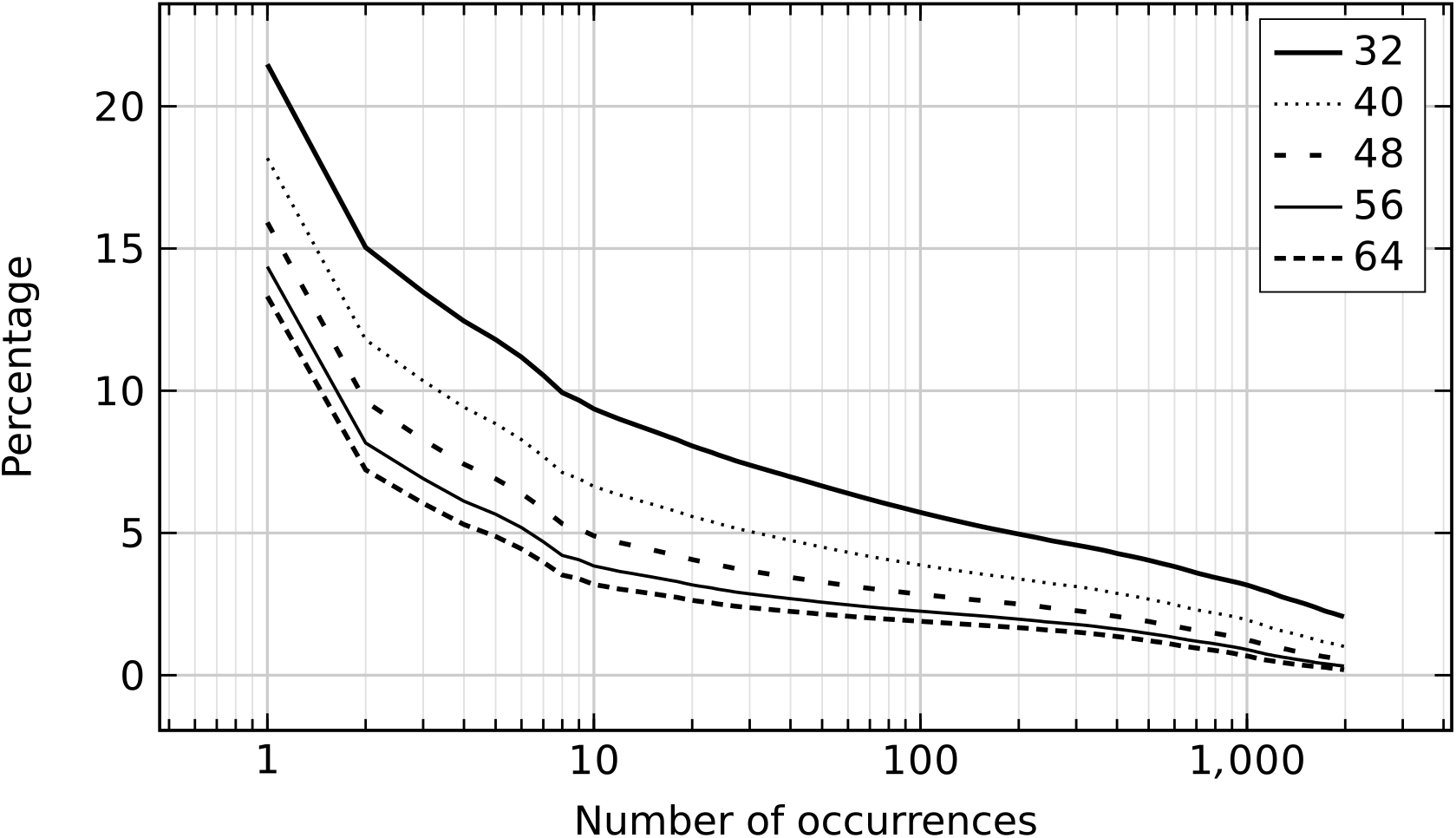
For a contiguous seed of a given weight (32, 40, 48, 56 and 64) we count the number of occurrences of same patterns within the human reference genome. Percentage of patterns (vertical axis) having more occurrences than a given occurrence (horizontal axis) is shown.

Secondly, for shorter reads, there is a chance to miss cases when a read has several substitutions. In [25] it is shown that a boolean sequence of *m* elements with at most *k* zeros contains at least one *l*-run of ones with 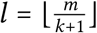. Therefore the use of 32-symbol substrings for 100 bp (150/200 bp) reads can guarantee to find all candidate positions if there are not more than 2 (3/5) substitutions. For 64-symbol substrings we get 0/1/2 substitutions, respectively.

In most practical applications majority of reads have complete matches or 1–2 substitutions. So, the use of long substrings (64 or more symbols) may allow to position reads correctly very quickly as the number of candidate positions is often small. Longer substrings may not work well when reads also contain insertions or deletions. So, there will be only a relative small number of reads without common substrings and those reads can be processed with shorter, e.g. 32-symbol, substrings.

### 2.3 Spaced seeds

Spaced seeds are good alternatives to contiguous chunks. Let there be two sequences 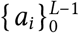 (a reference sequence) and 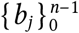 (a read). Suppose there is a boolean 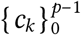 pattern/seed of length *p* ≤ *n* Consisting of ones and zeros. The number of ones is called the **weight** of the seed, while *p* is its **span** (or **length**). The Hamming distance for the reference subsequence starting at position *s* and the read subsequence starting at position *t* can be defined for 0 ≤ *s* ≤ *L* − *p* and 0 ≤ *t* ≤ *n* − *p* as

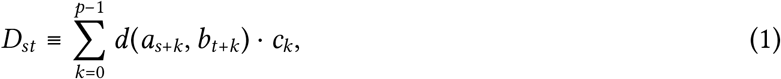

*d*(*A, B*) is the Hamming distance for two symbols *A* and *B*:

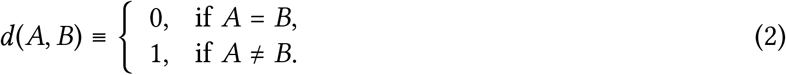

There may be four main types of output [14].

1. **Unique best hit**. Find *s* and *t* such as *D*_*st*_ is the smallest, report them if there is no other different pairs of *s* and *t*, otherwise report null.
2. **All valid hits**. Report all occurrences of *b* within *a*.
3. **Arbitrary hit**. For a given *δ* ≥ 0 report any pair of *s* and *t* such as *D*_*st*_ ≤ *δ*.
4. **All best hits**. Find the smallest value *β* of *D*_*st*_.If *β* ≤ *δ*, then report all pairs of *s* and *t*.

Let there be an arbitrary read 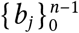 of length *n* containing no more than *m* substitutions. We may introduce a boolean array 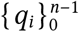 such that *q*_*i*_ = 0 if there is a substitution for the element *b*_*i*_ and 1, otherwise. Our goal is to find seeds 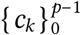 such that it is possible to translate them by *s* elements denote the seed as *c*^*s*^) and no 1-element of the translated seed meets 0-element of array *q*. A seed is designed successfully if for any permissible array *q* containing no more than *m* 0-elements we may find such translation *s*. Of course, the value of *s* depends on a given array *q*. Strict mathematical requirements can be written as: ∃*s* ∈ {0, 1, …, *n* − *p*}: ∀ *k* = {0, 1, …, *p* − 1} *q*_*s*+*k*_|*c*_*k*_= *q*_*s*+*k*_ (“|” is the bitwise OR). Both arrays *q* and *c* (and the translated array *c*^*s*^) can be padded with zeros. Then we expect that *q*|*c*^*s*^ *= q* for all elements.

### 2.4 Optimal remapping

Suppose we have found a spaced seed. For example, a seed shown in Figure 2. Its length is 59 and its weight is 32. Our aim is to form a contiguous array of ones with minimum number of remapping operations. If we can find such operations for 32-bit numbers, e.g. for **A**-component of the sequence, then we can form 128-bit instructions for all four components (**A, C, G, T**). We aim to use only 128-bit bitwise AND, OR or bit shift instructions. For the given example, the spaced seed has 15 gaps (where the seed has 0 values). So, if we want to form the contiguous seed by removing those gaps we need 14 shift operations. However, the order in which bits from the original seed are positioned in the contiguous seed is not important. Therefore, we may achieve this result by using only 3 shift operations. For convenience, we denote ones as letters A, B, C, D and all zeros are replaced by symbol “_”. All ones in the first 32 bits of the seed become A. Our goal is to shift the second 32 bits in such way that gaps/zeros of the first row can be filled with ones of the second row. Letters B, C, D are used to distinguish ones: letter B is used for the first shift (−8), letter C is for the second shift (−5) and letter D is for the third shift (15). Applying masking for the corresponding letters of the rows we get the following SIMD instructions to obtain a 128-bit **ACGT** structure corresponding to a contiguous seed.

**Figure 2.**
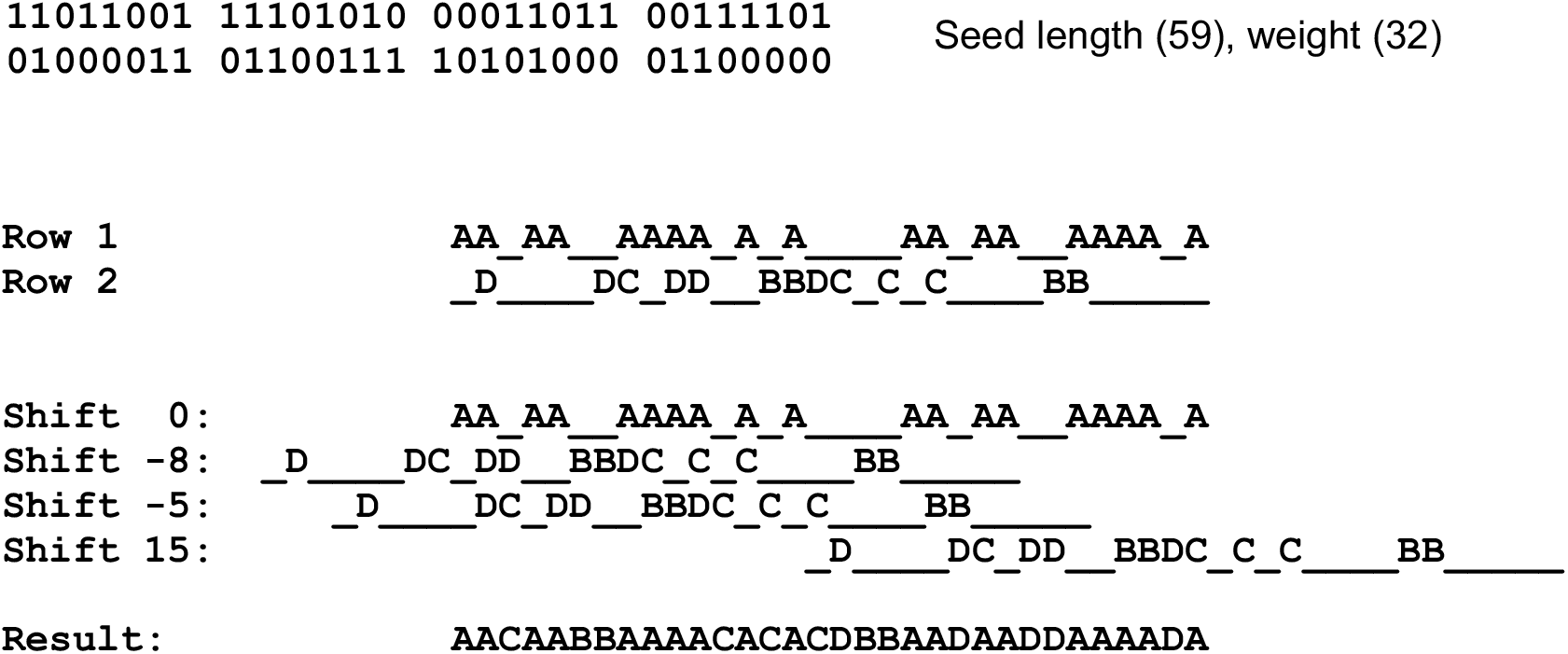
A possible procedure to form a contiguous seed for a spaced seed of length 59 and weight 32. For better presentation it is split into two rows (32 bit each) with extra gaps for each 8 elements.

~~~
**__m128i c, t, s;
c = _mm_set1_epi32(0xbcd8579b);
res[0] = _mm_and_si128(m[0], c);
c = _mm_set1_epi32(0×06006000);
t = _mm_and_si128(m[1], c);
s = _mm_srli_epi32(t, 8);
res[0] = _mm_or_si128(res[0], s); c = _mm_set1_epi32(0×00150080);
t = _mm_and_si128(m[1], c); s = _mm_srli_epi32(t, 5);
res[0] = _mm_or_si128(res[0], s); c = _mm_set1_epi32(0×00008642);
t = _mm_and_si128(m[1], c);
s = _mm_slli_epi32(t, 15);
res[0] = _mm_or_si128(res[0], s);**
~~~

There may be several possible combinations to form contiguous (even for a given number of shifts). Similar ideas can be used to form seeds of other lengths/weights.

We explain how the problem can be solved for the given example (weight 32 and two rows), with more details in Supplementary Material S4. The number of zeros for the first row is 13. We shift the second row (from −31 to 31) and for each shift value we check if any ones from the second row can be at same positions as zeros of the first row. If the shift provides us with such a chance, then we form a vector. The length of the vector is the number of zero elements for the first row of the seed. Each element of the vector is 0 if there is no 1-element of the shifted second row, otherwise it is the index of this element. As the result we get 45 such vectors and can form a (45 × 13)-matrix. For a given number *k* of shifts we need to pick up *k* rows of this matrix, apply a reduction procedure to be sure the matrix is a valid one:

1. all 1-indexes of the second row of the seed are present in the matrix;
2. there are no rows/columns of zeros;
3. if a non-zero element is the only non-zero element in a column, then for all other columns the same non-zero elements are set to 0;
4. if there are several non-zero elements in a column and one of non-zero elements is the only element in the matrix, then all other elements of the column can be set to zero.

If after the reduction procedure there are still several same non-zero elements, then for a column with *m* > 1 non-zero elements we form *m* matrices which are the same as the original matrix but only one non-zero element in the column. For each of those *m* matrices we apply the above reduction procedure. After the procedure we get one or several matrices with all columns containing only one non-zero element and each non-zero element is unique in the matrix. Those matrices allow us to form masks for 1-elements of the seed’s row and find shift values.

## 3 Results

### 3.1 General properties of spaced seeds

Let us consider all reads of length *n*, containing *k* substitutions. We want to find all spaced seeds of length *m* and weight *w* that can be used for any read of the given class. We assume that each seed starts and ends with 1.. Denote all possible seeds as *S* (*n, k, m, w*). If there are no such seeds, then the set is the empty set ∅.

1. ∀ *ξ* ∈ *S* (*n, k, m, w*), ∀ *p* > *n*: *ξ* ∈ *S* (*p, k, m, w*), i.e. if a seed is valid for reads of a given length, then it is also valid for longer reads. Therefore we have *S*(*n, k, m, w*) ⊂ *S*(*n* + 1, *k, m, w*).
2. ∀ *ξ* ∈ *S* (*n, k, m, w*), ∀ *r* < *k*: *ξ* ∈ *S* (*n, r, m, w*), i.e. if a seed is valid for a given number of substitutions, then it is also valid for cases of less substitutions. We have *S*(*n, k, m, w*) ⊂ *S*(*n,k* − 1,, *m, w*).

To check if a seed *ξ* ∈ *S* (*n, k, m, w*), w is valid we need to generate all possible reads of length *n* with *k* substitutions or, equivalently, a binary *n*-array with *k* zeros. There are 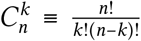 possible reads. For each read we need to translate the seed. As the read has length *m* we may perform (*n* − *m* + 1) translations.

In Figure 3 we see all spaced seeds found for *n* =11 and *k* =2. The seeds are combined in groups depending on the value of weight *w*. For any valid seed of weight w its sub-seeds (obtained by either reducing its length or setting some bits to zero but having 1s as its first/last elements) should also be valid seeds. For example, `11101` has weight of 4, all its sub-seeds of weight 3 are `111`, `11001`, `10101` and `1101` are also present in Figure 3. Therefore, we may construct all seeds iteratively. We start with all seeds of length 1, i.e. `1`, then continue with longer seeds by extending seeds found on previous steps and checking if the necessary condition (all sub-seeds are already in the list of found seeds). If a found seed is of length *p* and a candidate seed should of length *m*, then we pad the found seed by a contiguous (*m*− *p* – 1) -array of zeros and one 1-element.

**Figure 3.**
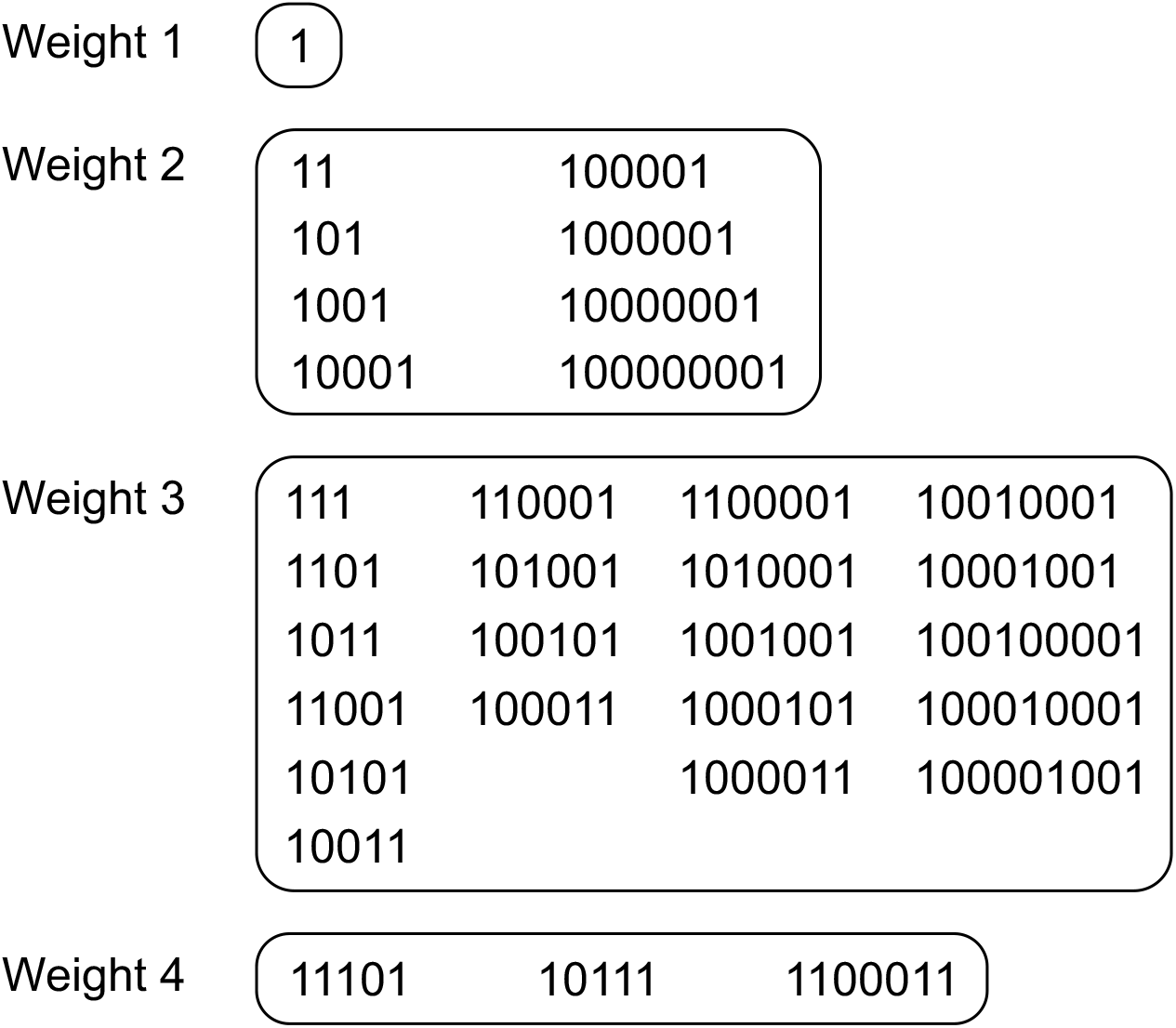
All possible patterns for a read of length 11 and 2 substitutions. The patterns are grouped based on a number of non-zero elements.

### 3.2 Periodic spaced seeds

For a given length *n* of a read and number *k* of substitutions there may be many valid seeds. We should agree how to select best of them. Our aim is to use seeds for sequence alignment problems. Therefore the best option is to choose seeds of the maximum weight. In this case we reduce a number of candidate positions within a reference sequence. When a seed is designed, we have no information about a read. So, the chance to find a subsequence of weight w should be 4 times more likely than for a subsequence of weight (*w* + 1). Therefore, when there is a set of possible seeds we choose seeds of maximum weight. For the case shown in Figure 3 we have three seeds (`11101`, `10111` and `1100011`).

For a human genome some subsequences may appear in several places. With longer sequences there is a chance that a frequent sequence will also contain unique patterns. So, among seeds of maximum weight we choose seeds of maximum length. For the case discussed above the best seed is `1100011`. Examples of other spaced seeds for *k* = 3 substitutions are shown in Figure 4.

**Figure 4.**
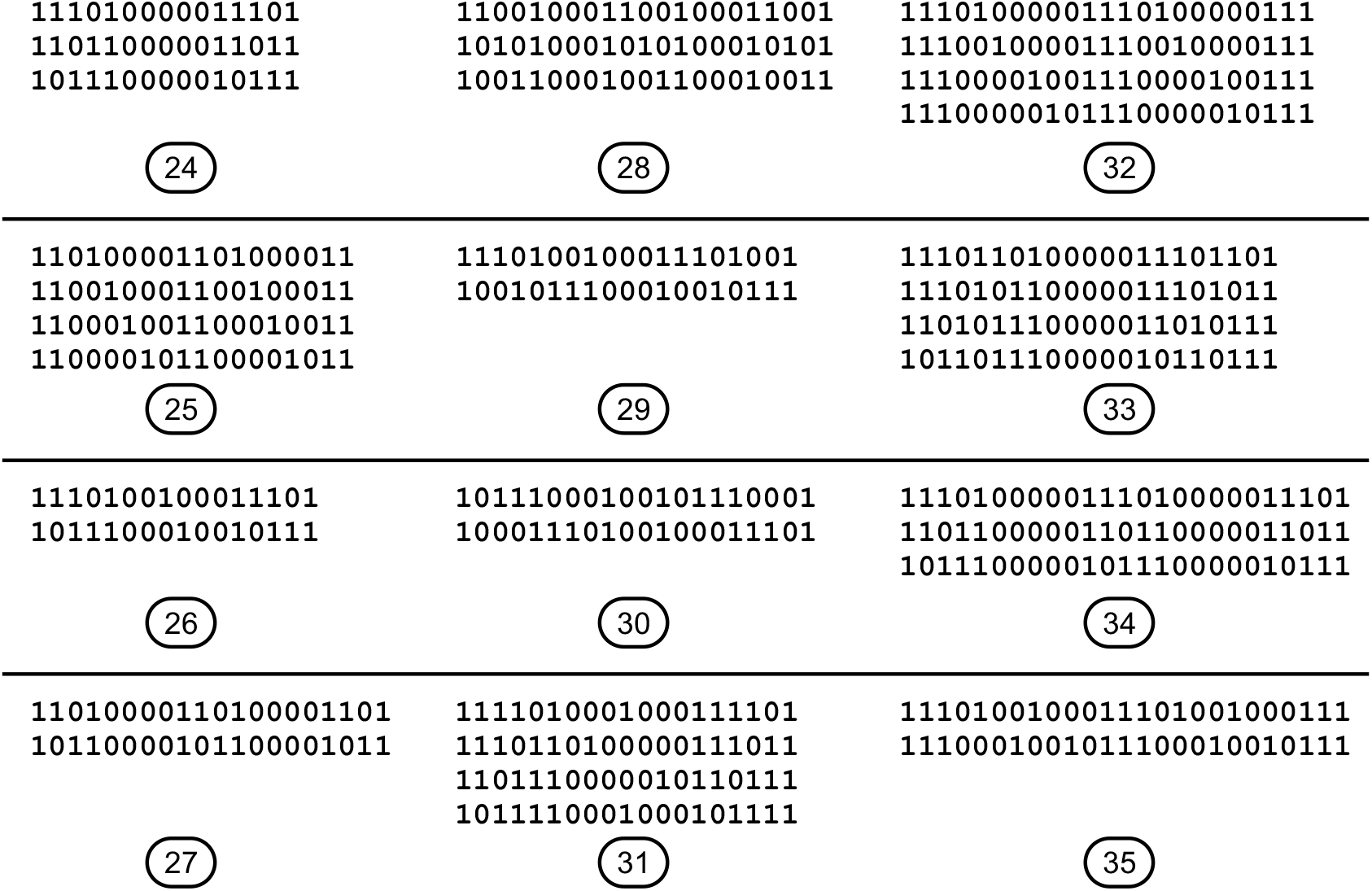
For a given length (numbers in ovals) of reads patterns with a maximum number of non-zero elements are found, 3 substitutions. Only maximum-length patterns are shown.

We may see that “maximum weight/maximum length” seeds are periodic patterns. For the examples provided we may find a shorter boolean pattern such that the spaced seed is a concatenation of several same patterns. In Figure 5 the corresponding shorter patterns are shown. A spaced seed usually contains several whole patterns and a subpattern ending with `1`. This is a purely practical observation (without any proof). There is always a chance that for long reads best spaced seeds may have a different structure. However, in order to avoid computationally expensive procedures we suppose that best seeds have periodic structures.

**Figure 5.**
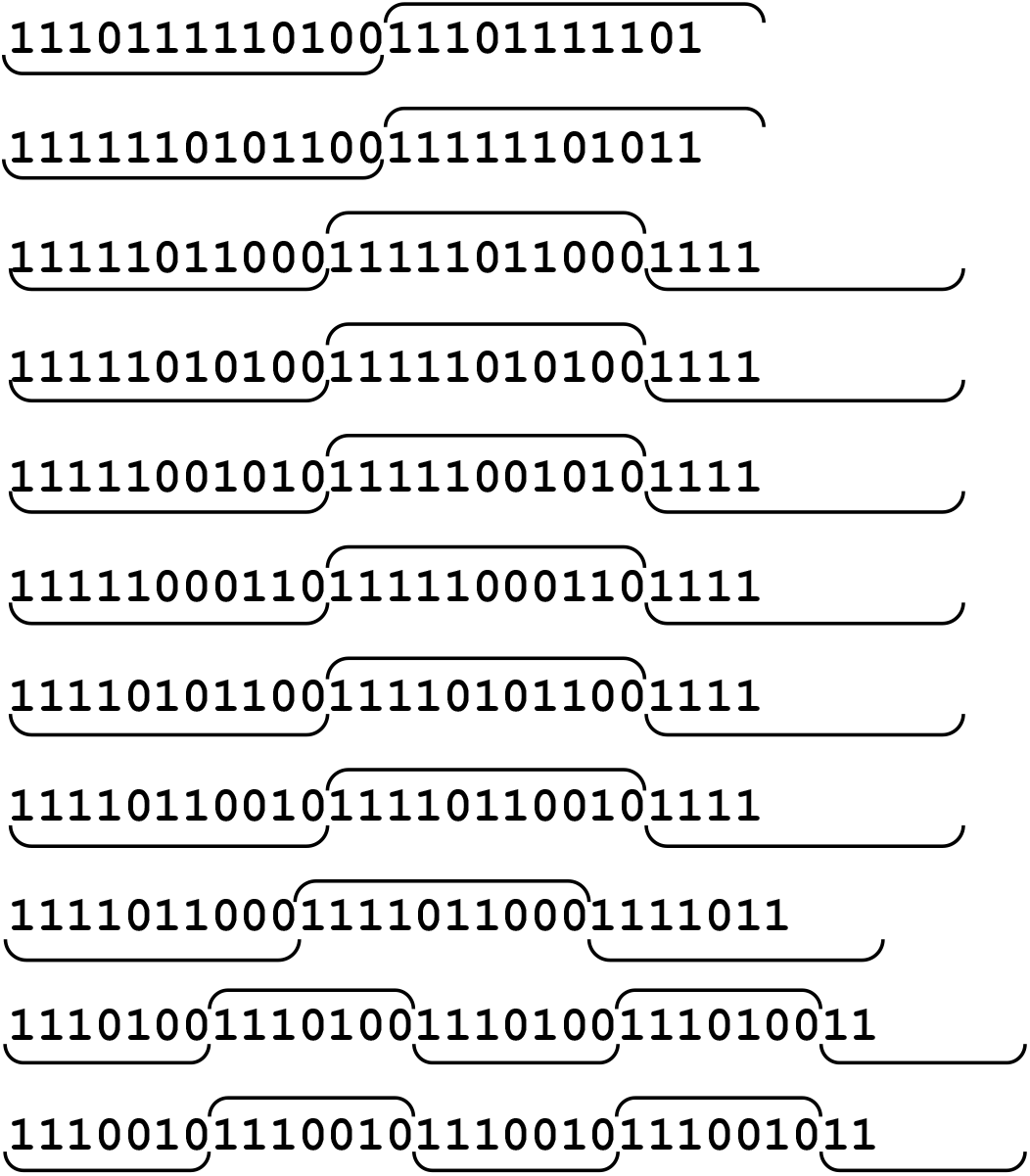
All maximum-length patterns found for a read of length 36 and 2 substitutions. Flipped patterns are not shown. The patterns are periodic, the corresponding sub-patterns are indicated.

### 3.3 Numerical procedure

For a given length *n* of a read, weight *w* and period *T* (the length of a periodic structure starting with 1 and ending with 0) a code to generate all possible spaced seeds is written. For each candidate periodic seed we randomly generate positions of substitutions within a read (millions of various combinations). Fixing weight *w*, number of substitutions *k* and the period *T* we vary the length n of reads. According to general properties of spaced seeds, by reducing the length we reduce the number of spaced seeds. In Figure 6 we see how the number of minimum lengths of reads are found as a function of period *T*. Computational complexity is increased with the number of substitutions and length of a read. Therefore the maximum periods were 25–28 depending on the number of substitutions (Supplementary Material S5). The best seeds are chosen from seeds permitting smallest values of reads’ lengths (see Table 2 and Supplementary Material S6).

**Table 2.**
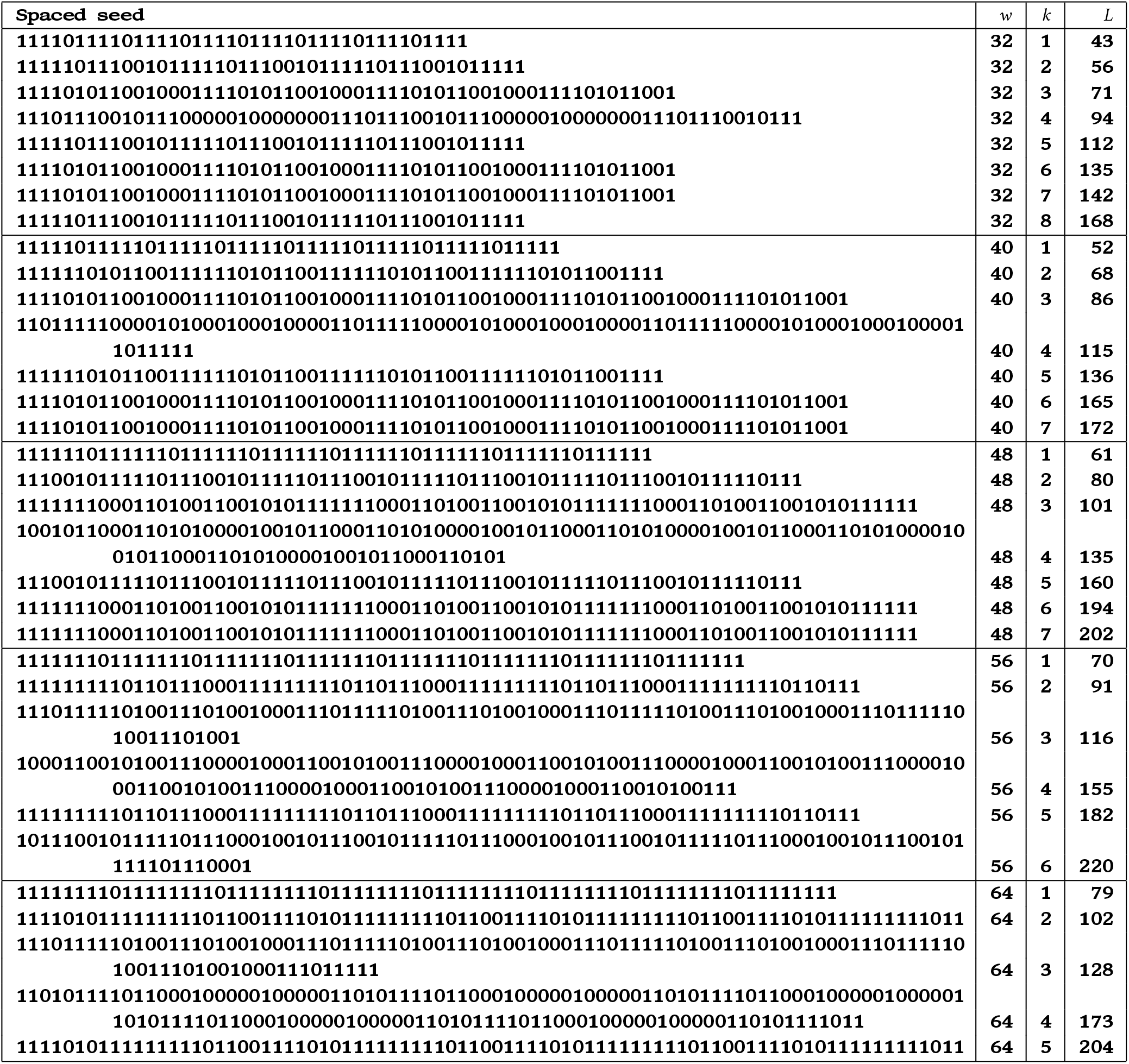
Spaced seeds of weight *w* that can be used for reads of at least a given length *L* with no more than *k* substitutions.

**Figure 6.**
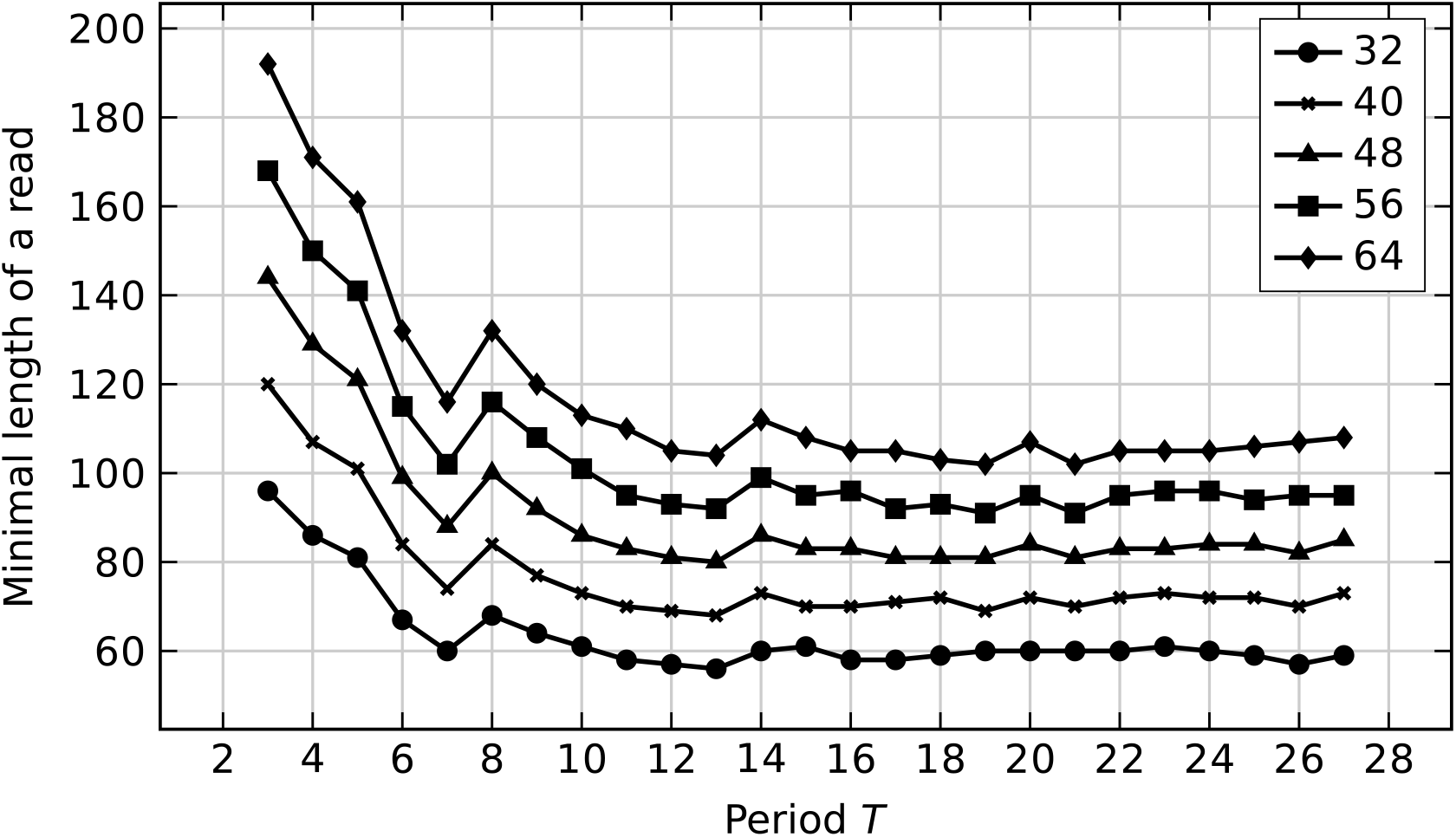
The minimal value of reads’ lengths required for periodic spaced seeds of given weights (up to 2 substitutions are allowed).

**Figure 7.**
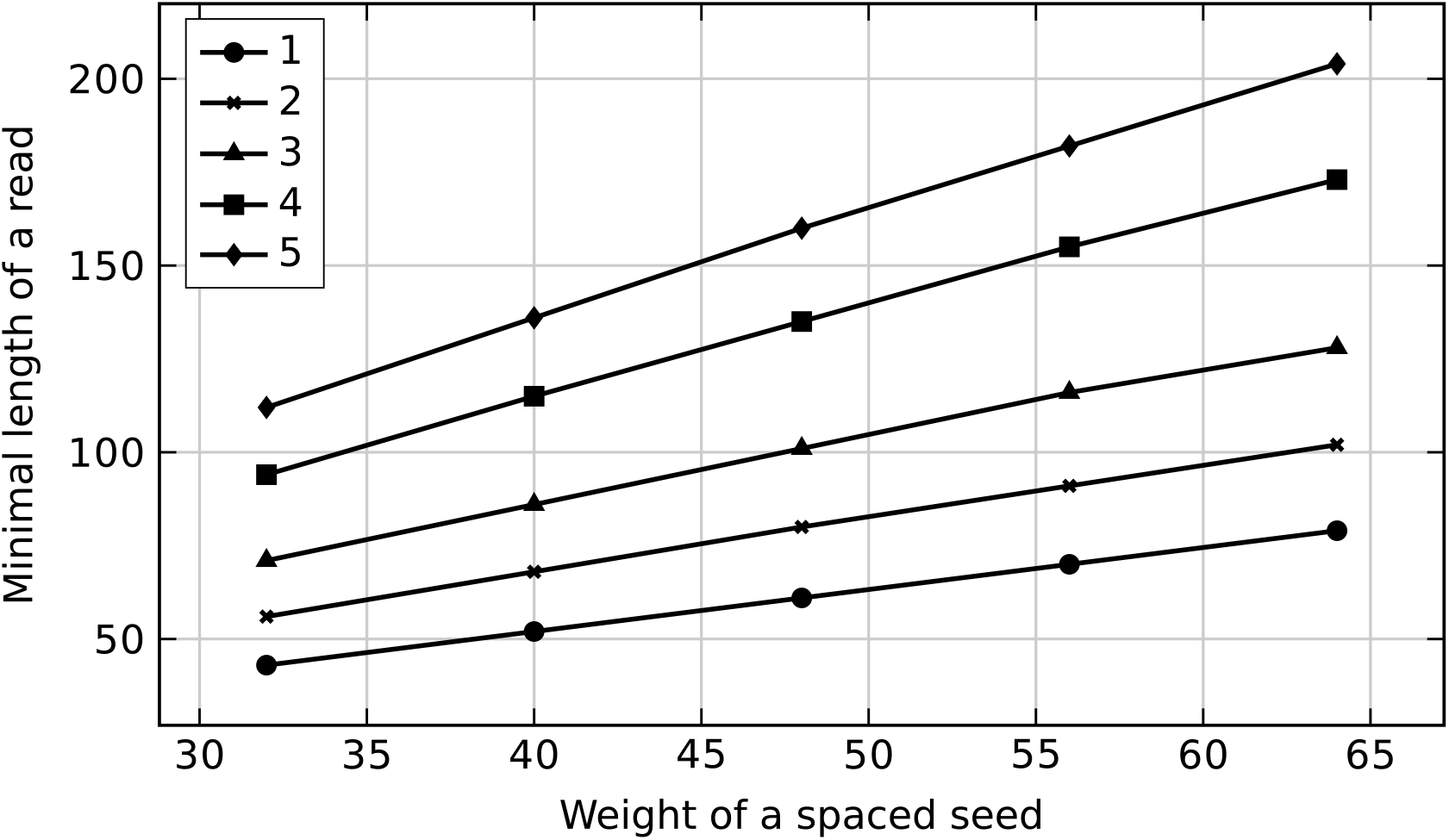
Minimum values for a read’s length as a function of weight of spaced seeds found for different number of substitutions (1, 2, 3, 4, 5).

For each best periodic seed we identify optimal remapping procedure to convert sequences with gaps into “signature” numbers by using as less SIMD procedures as possible. The output results of the software for the best seeds can be found in Supplementary Material S7.

## 4 Discussion

Software tools used to generate results of the paper are available at https://github.com/vtman/VSTseed. The authors worked only with seeds of specific weights (32, 40, 48, 56 and 64) and periods (usually less than 29). In principle, other weights, e.g. multiples of 4, can be used. In this case each pair (“position”, “signature”) will require an integer number of bytes. Other periods can also be considered with possibly other seeds to be found.

The minimum length of reads required for seeds of given weight is almost a linear function (or more strictly, an affine function) of a number of substitutions allowed. As the minimum length of a read for a contiguous seed of weight *w* can be written as (*k* + 1) *w* + *k*, the corresponding lengths for the best spaced seeds are near 53% of (*k* + 1) *w* + *k*. For example, contiguous seeds of weight *w* =48 and at most *k* =3 substitutions require lengths of at least 195 bp, but corresponding spaced seeds must be at least 101 bp, i.e. 52%.

There is usually a very small number of seeds valid for the reads of minimal lengths. However, when reads’ lengths are given, e.g. *L =*120, and weight of seeds is chosen, e.g. *w* =48, there may be many seeds to be used that guarantee full sensitivity. For the known *L* and *w* values we obtain full sensitivity for the case of at most 3 substitutions. We cannot use *k* =4 as the minimum requirement is *L* =135, however the minimum requirement for *k* =3 is *L* =101, so more seeds are available. Some of those seeds might have shorter conversion “sequence”–”signature” procedures (less SIMD instructions).

Weight of a seed is the main parameter determining the number of candidate positions within a reference sequence. Percentage of patterns determined by different seeds of the same weight that have a given number of occurrences within the human reference genome vary slightly. The curves found for contiguous seeds and shown in Figure 1 are similar to curves obtained for spaced seed. Curves for lengthier seeds usually below the curves for contiguous seeds but within [0, 0.3%] interval.

When a spaced seed and corresponding SIMD “signature” instructions are found, indexing of the whole genome is a relatively fast procedure for a modern budget computer. For example, a PC with Intel I5-10600KF CPU (6 cores, 4.1 GHz) and NMVe storage (2400/1950 MBps read/write speed) form unsorted index files for *w*= 32 within 28.6 s (31.8 GB of data) and for *w* = 64 within 92.7 s (54.9 GB). To sort the files 75.2 s and 115.8 s are needed. The index files should be generated only once when reads’ lengths or seeds are changed.

For read alignment problems it may be reasonable to generate index files for several weights. Usually aligned reads have a very small number of substitutions with respect to a reference sequence. Therefore, we may consider seeds of greater weights, check all candidate positions for each read and only in case of large values of Hamming distance between a read and a chunk of the reference sequence we attempt to find other candidate positions using seeds of smaller weight. For example, if our reads have length 140, then we consider seeds of weight 64. In this case we are guaranteed to find all locations when the number of substitutions is no more than 3. Once all reads are processed, we consider reads with no candidate locations found, e.g. when there is no corresponding positions in the reference sequence for a given “signature”, and reads with higher values of Hamming distance. For these reads we may use seeds of weight 32 allowing us to have up to 6 substitutions.

An optimal seed found for reads of length *L*, weight *w* and at most *k* substitutions will help us to pick up some positions in case of insertions or deletions. The shorter length of a seed is, the higher chance to find those positions is. Of course, in case of very small number of indels we may generate corresponding signatures directly from a given read. If *L*_*s*_ is a length of a seed, then in case of no indels we should generate (*L*− *L* _*s*_ + 1) “signatures” for each read. For a single insertion we obtain 4(*L* + 1) new reads of length (*L*+ 1). Thus may obtain up to 4(*L*+ 1)(*L*− *L*_*s*_ + 2) “signatures”. However, the real number of “signatures” is smaller, since several new reads have same patterns.

Shorter seeds tend to have slightly larger number of candidate positions for each seed. As the result, shorter seeds will force us to consider more candidate positions. The authors plan to discuss strategies to choose periodic seeds, combine several ones for faster mapping and their embedding in a new alignment software in future papers.

