## Supplementary material for "VSTseed: periodic spaced seeds for reads with substitutions"

#### S1. Digital representation of sequences

Each symbol of a sequence is either an element of alphabet $\Sigma = \{A, C, G, T\}$ or symbol $N$. Thus we need 5 numbers $(0, 1, 2, 3, 4)$ or 3 bits to code a symbol.

| | | | | | | | | | | | | | | | |
|---|---|---|---|---|---|---|---|---|---|---|---|---|---|---|---|
| *A* | 000 | *C* | 001 | *G* | 010 | *T* | 011 | *N* | 100 | — | 101 | — | 110 | — | 111 |

For two symbols we need 5 bits to code them.

| | | | | | | | | | | | | | | | |
|---|---|---|---|---|---|---|---|---|---|---|---|---|---|---|---|
| *AA* | 00000 | *AC* | 00001 | *AG* | 00010 | *AT* | 00011 | *AN* | 00100 | *CA* | 00101 | *CC* | 00110 | *CG* | 00111 |
| *CT* | 01000 | *CN* | 01001 | *GA* | 01010 | *GC* | 01011 | *GG* | 01100 | *GT* | 01101 | *GN* | 01110 | *TA* | 01111 |
| *TC* | 10000 | *TG* | 10001 | *TT* | 10010 | *TN* | 10011 | *NA* | 10100 | *NC* | 10101 | *NG* | 10110 | *NT* | 10111 |
| *NN* | 11000 | — | 11001 | — | 11010 | — | 11011 | — | 11100 | — | 11101 | — | 11110 | — | 11111 |

And for three symbols 7 bits are required.

| | | | | | | | | | | | |
|---|---|---|---|---|---|---|---|---|---|---|---|
| *AAA* | 0000000 | *AAC* | 0000001 | *AAG* | 0000010 | *AAT* | 0000011 | *AAN* | 0000100 | *ACA* | 0000101 |
| *ACC* | 0000110 | *ACG* | 0000111 | *ACT* | 0001000 | *ACN* | 0001001 | *AGA* | 0001010 | *AGC* | 0001011 |
| *AGG* | 0001100 | *AGT* | 0001101 | *AGN* | 0001110 | *ATA* | 0001111 | *ATC* | 0010000 | *ATG* | 0010001 |
| *ATT* | 0010010 | *ATN* | 0010011 | *ANA* | 0010100 | *ANC* | 0010101 | *ANG* | 0010110 | *ANT* | 0010111 |
| *ANN* | 0011000 | *CAA* | 0011001 | *CAC* | 0011010 | *CAG* | 0011011 | *CAT* | 0011100 | *CAN* | 0011101 |
| *CCA* | 0011110 | *CCC* | 0011111 | *CCG* | 0100000 | *CCT* | 0100001 | *CCN* | 0100010 | *CGA* | 0100011 |
| *CGC* | 0100100 | *CGG* | 0100101 | *CGT* | 0100110 | *CGN* | 0100111 | *CTA* | 0101000 | *CTC* | 0101001 |
| *CTG* | 0101010 | *CTT* | 0101011 | *CTN* | 0101100 | *CNA* | 0101101 | *CNC* | 0101110 | *CNG* | 0101111 |
| *CNT* | 0110000 | *CNN* | 0110001 | *GAA* | 0110010 | *GAC* | 0110011 | *GAG* | 0110100 | *GAT* | 0110101 |
| *GAN* | 0110110 | *GCA* | 0110111 | *GCC* | 0111000 | *GCG* | 0111001 | *GCT* | 0111010 | *GCN* | 0111011 |
| *GGA* | 0111100 | *GGC* | 0111101 | *GGG* | 0111110 | *GGT* | 0111111 | *GGN* | 1000000 | *GTA* | 1000001 |
| *GTC* | 1000010 | *GTG* | 1000011 | *GTT* | 1000100 | *GTN* | 1000101 | *GNA* | 1000110 | *GNC* | 1000111 |
| *GNG* | 1001000 | *GNT* | 1001001 | *GNN* | 1001010 | *TAA* | 1001011 | *TAC* | 1001100 | *TAG* | 1001101 |
| *TAT* | 1001110 | *TAN* | 1001111 | *TCA* | 1010000 | *TCC* | 1010001 | *TCG* | 1010010 | *TCT* | 1010011 |
| *TCN* | 1010100 | *TGA* | 1010101 | *TGC* | 1010110 | *TGG* | 1010111 | *TGT* | 1011000 | *TGN* | 1011001 |
| *TTA* | 1011010 | *TTC* | 1011011 | *TTG* | 1011100 | *TTT* | 1011101 | *TTN* | 1011110 | *TNA* | 1011111 |
| *TNC* | 1100000 | *TNG* | 1100001 | *TNT* | 1100010 | *TNN* | 1100011 | *NAA* | 1100100 | *NAC* | 1100101 |
| *NAG* | 1100110 | *NAT* | 1100111 | *NAN* | 1101000 | *NCA* | 1101001 | *NCC* | 1101010 | *NCG* | 1101011 |

| | | | | | | | | | | | |
|---|---|---|---|---|---|---|---|---|---|---|---|
| *NCT* | 1101100 | *NCN* | 1101101 | *NGA* | 1101110 | *NGC* | 1101111 | *NGG* | 1110000 | *NGT* | 1110001 |
| *NGN* | 1110010 | *NTA* | 1110011 | *NTC* | 1110100 | *NTG* | 1110101 | *NTT* | 1110110 | *NTN* | 1110111 |
| *NNA* | 1111000 | *NNC* | 1111001 | *NNG* | 1111010 | *NNT* | 1111011 | *NNN* | 1111100 | — | 1111101 |
| — | 1111110 | — | 1111111 | | | | | | | | |

#### S2. Intel SIMD instructions

Intel intrinsics instructions (https://software.intel.com/sites/landingpage/IntrinsicsGuide/).

We provide a list of intrinsics useful for generation of optimal spaced seeds. Performance numbers for Skylake architecture are shown in the table.

| **Name** | **Description** | **Latency** | **Throughput (CPI)** |
|---|---|---|---|
| `__m128i _mm_load_si128 (__m128i const* mem_addr)` | Load 128-bits of integer data from memory into dst. | 6 | 0.5 |
| `void _mm_store_si128 (__m128i* mem_addr, __m128i a)` | Store 128-bits of integer data from a into memory. | 5 | 1 |
| `__m128i _mm_set_epi32 (int e3, int e2, int e1, int e0)` | Set packed 32-bit integers in dst with the supplied values. | | |
| `__m128i _mm_set1_epi32(int a)` | Broadcast 32-bit integer a to all elements of dst. | | |
| `__m128i _mm_or_si128 (__m128i a, __m128i b)` | Compute the bitwise OR of 128 bits (representing integer data) in a and b, and store the result in dst. | 1 | 0.33 |
| `__m128i _mm_and_si128 (__m128i a, __m128i b)` | Compute the bitwise AND of 128 bits (representing integer data) in a and b, and store the result in dst. | 1 | 0.33 |
| `int _popcnt32 (int a)` | Count the number of bits set to 1 in 32-bit integer a. | 3 | 1 |
| `__m128i _mm_shuffle_epi8 (__m128i a, __m128i b)` | Shuffle packed 8-bit integers in a according to shuffle control mask in the corresponding 8-bit element of b. | 1 | 1 |
| `__m128i _mm_slli_epi32 (__m128i a, int imm8)` | Shift packed 32-bit integers in a left by imm8 while shifting in zeros. | 1 | 0.5 |
| `__m128i _mm_srli_epi32 (__m128i a, int imm8)` | Shift packed 32-bit integers in a right by imm8 while shifting in zeros. | 1 | 0.5 |

#### S3. SIMD representation of sequences

Each 32-symbol sequence we write as a 128-bit number. An $i$-th bit from the first 32 bits is 1 if the $i$-th symbol in the sequence is $A$, otherwise it is 0. The next 32 bits are for symbol $C$, then for symbol $G$ and $T$ respectively. If there is a symbol $N$, then all bits in $A$, $C$, $G$ and $T$ arrays are 0.

Let there be a 64-symbol sequence. We may split it into two 32-symbol sequences (`m1` and `m2`). Suppose we also select a 32-symbol sequence `m3` (starting at 14th element of the original sequence).

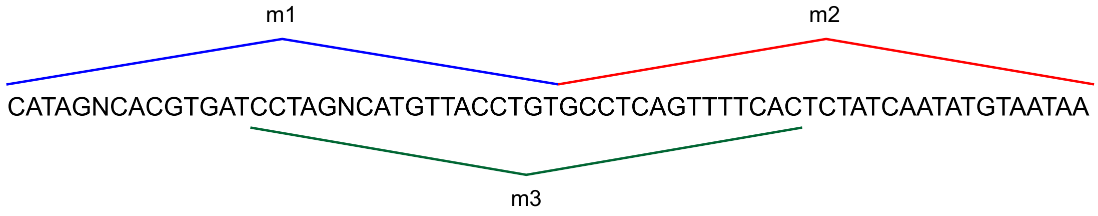


All $A$, $C$, $G$ and $T$ components are shown below. Note that if there is no $N$ symbol, then after applying bitwise OR operations to $A$, $C$, $G$ and $T$ components we get `0xffffffff`.

| m1 | CATAGNCACGTGATCCTAGNCATGTTACCTGT | |
|---|---|---|
| $A$ | 01010001000010000100010000100000 | 0x0422108a |
| $C$ | 10000010100000011000010000000011000 | 0x1810c141 |
| $G$ | 00001000001010000000100001000000010 | 0x40840a10 |
| $T$ | 0010000000010010010000001011000101 | 0xa3412404 |
| $A\|C\|G\|T$ | 11111011111111111111011111111111 | 0xfff7ffdf |

| m2 | GCCTCAGTTTTCACTCTATCAATATGTAATAA | |
|---|---|---|
| $A$ | 000001000000010000100110100011011 | 0xd8b21020 |
| $C$ | 01101000000101010001000000000000 | 0x0008a816 |
| $G$ | 10000010000000000000000000001000000 | 0x02000041 |
| $T$ | 00010001111000101010001010100100 | 0x25454788 |
| $A\|C\|G\|T$ | 11111111111111111111111111111111 | 0xffffffff |

| m3 | CCTAGNCATGTTACCTGTGCCTCAGTTTTCAC | |
|---|---|---|
| $A$ | 00010001000010000000000001000000010 | 0x40801088 |
| $C$ | 11000001000000011000001101000000101 | 0xa0586043 |
| $G$ | 00001000010000000101000000100000000 | 0x01050210 |
| $T$ | 001000000101100010100010001111000 | 0x1e228d04 |
| $A\|C\|G\|T$ | 11111011111111111111111111111111 | 0xffffffdf |

All three structure can be set using SIMD instructions.

```
m1 = _mm_set_epi32 (0xa3412404, 0x40840a10, 0x1810c141, 0x0422108a);
m2 = _mm_set_epi32 (0x25454788, 0x02000041, 0x0008a816, 0xd8b21020);
m3 = _mm_set_epi32 (0x1e228d04, 0x01050210, 0xa0586043, 0x40801088);
```

A long sequence can be split into 32-symbol chunks which we write down as 128-bit structures. We need to store only structures obtained with 32-symbol steps. All intermediate structures (like `m3`) can be found with shift and bitwise OR operations (at most 3 SIMD instructions). For example,

```
__m128i m1, m2, m3, m4, m5, m6;
m4 = _mm_srli_epi32(m1, 14);
m5 = _mm_slli_epi32(m2, 32-14);
```

```
m6 = _mm_or_si128(m4, m5);
// m1 = {0x0422108a, 0x1810c141, 0x40840a10, 0xa3412404}
// m2 = {0xd8b21020, 0x0008a816, 0x02000041, 0x25454788}
// m4 = {0x00001088, 0x00006043, 0x00010210, 0x00028d04}
// m5 = {0x40800000, 0xa0580000, 0x01040000, 0x1e200000}
// m6 = {0x40801088, 0xa0586043, 0x01050210, 0x1e228d04}
// m3 = m6
```

We count the number of same symbols (except $N$) for `m1` and `m2`

```
unsigned int ures;
int icount;
__m128i m1, m2, m3, m4, m5, m6, m7;
m3 = _mm_and_si128(m1, m2);
m4 = _mm_bsrli_si128(m3, 8);
m5 = _mm_or_si128(m3, m4);
m6 = _mm_bsrli_si128(m5, 4);
m7 = _mm_or_si128(m5, m6);
ures = _mm_extract_epi32(m7, 0);
icount = _mm_popcnt_u32(ures);
// m3 = {0x00221000, 0x00008000, 0x00000000, 0x21410400}
// m4 = {0x00000000, 0x21410400, 0x00000000, 0x00000000}
// m5 = {0x00221000, 0x21418400, 0x00000000, 0x21410400}
// m6 = {0x21418400, 0x00000000, 0x21410400, 0x00000000}
// m7 = {0x21639400, 0x21418400, 0x21410400, 0x21410400}
// ures = 0x21639400
// icount = 9
```

| | | *A* | *C* | *G* | *T* |
|---|---|---|---|---|---|
| m1 | CATAGNCACGTGATCCTAGNCATGTTACCTGT | 0x0422108a | 0x1810c141 | 0x40840a10 | 0xa3412404 |
| m2 | GCCTCAGTTTTCACTCTATCAATATGTAATAA | 0xd8b21020 | 0x0008a816 | 0x02000041 | 0x25454788 |
| m1 & m2 | __________T_A__CTA___AT_T____T__ | 0x00221000 | 0x00008000 | 0x00000000 | 0x21410400 |
| 0x00221000 OR 0x00008000 OR 0x00000000 OR 0x21410400 = 0x21639400 | | | | | |
| _mm_popcnt_u32(0x21639400) = 9 | | | | | |

### S4. Converting spaced seeds to contiguous arrays

Suppose we have found a spaced seed. For example,

$$11011001111010100001101100111101010000110110011110101000011 \quad (1)$$

The length of the seed is 59 and its weight (number of 1s) is 32. We want to rearrange indices of the original seed and form a contiguous pattern of length/weight 32. The simplest approach is just remove all zero elements and preserve the order of 1s like below.

```
11011001111010100001101100111101010000110110011110101000011
```

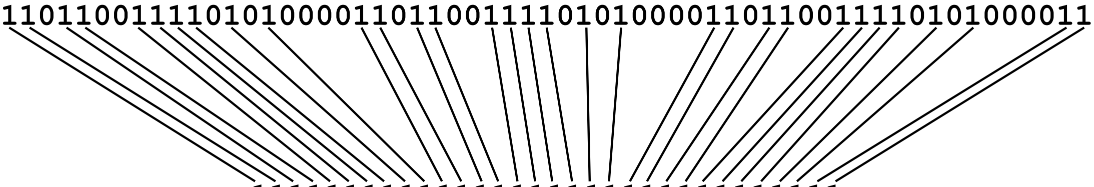

```
11111111111111111111111111111111
```

However, for the given spaced seed we will need at least 14 shift operations as there are 15 contiguous chunks of zeros.

We aim to use 128-bit SIMD instructions applied to *ACGT*-sequences (first 32 bits are for *A* component, next 32 bits are for *C*, etc.). So, we pad the original seed with zeros, so the new seed length is a multiple of 32 and split it into 32-bit arrays (each array corresponds to a new row):

```
row 1:  11011001111010100001101100111101
row 2:  01000011011001111010100001100000
```

We may try to solve the problem by the following approach. All 1s of the first row do not change their position, we translate the second row with respect to the first one and try to find such shifts, so 1s of the second row are just below 0s of the first row. We need to perform multiple shifts. One possible combination of shifts is shown below

```
                      11011001111010100001101100111101
shift  -8:    01000011011001111010100001100000
shift  -5:       01000011011001111010100001100000
shift  15:                                   01000011011001111010100001100000
```

After masking has been applied to the second row we have

```
                      11011001111010100001101100111101
shift  -8:    0 0000   0   0011   0 0 00001100000
shift  -5:       0 0000 10  00    101010000  00000
shift  15:                                   0100001 01100  1 0 0 0000  00000
```

Then we get the 32-bit contiguous array

```
11111111111111111111111111111111
```

For shortness 32-bit masking can be written as hex numbers (where `0x00000001` corresponds to a mask of all zeros except the left element which is one), so

| row | shift | ≪ or ≫ | value | mask, hex | mask, binary |
|---|---|---|---|---|---|
| 1 | 0 | — | 0 | `0xbcd8579b` | `11011001111010100001101100111101` |
| 2 | −8 | ≫ | 8 | `0x06006000` | `00000000000001100000000000001100000` |
| 2 | −5 | ≫ | 5 | `0x00150080` | `00000001000000000101010000000000000` |
| 2 | 15 | ≪ | 15 | `0x00008642` | `01000010011000010000000000000000` |

If we apply corresponding shift to hex values of masks, perform OR instructions for the resultant values, then we should get `0xffffffff`.

| original mask | shift | resultant mask, hex | resultant mask, binary |
|---|---|---|---|
| `0xbcd8579b` | — | `0xbcd8579b` | `11011001111010100001101100111101` |
| `0x06006000` | ≫ 8 | `0x00060060` | `00000110000000000011000000000000000` |
| `0x00150080` | ≫ 5 | `0x0000a804` | `00100000000010101000000000000000000` |
| `0x00008642` | ≪ 15 | `0x43210000` | `00000000000000000100001001100010` |
| Total | | `0xffffffff` | `11111111111111111111111111111111` |

The mapping is shown below

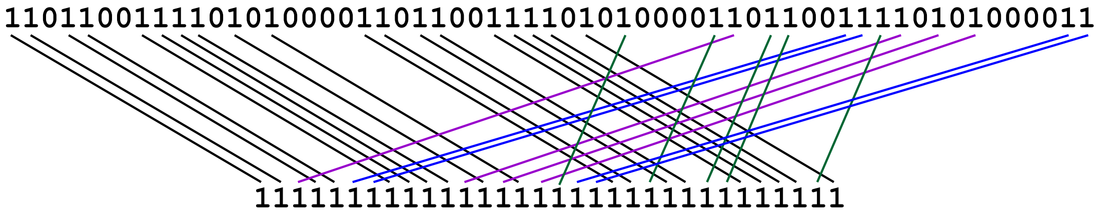


Now we discuss how to find possible shifts that will provide us with a solution. There are 13 zeros in the first row and 32 elements in each row. The first row is fixed and we just shift the second row and pad it with zeros if it is needed. Shift values can vary from $-31$ to 31. For each shift we check if any 1s from the second row are at same positions as 0s of the first row. In case of success we write a vector of length 13 and write corresponding indexes of 1s, otherwise we write 0. If there are no such elements, then we consider another shift value. In this way we create a $45 \times 13$ matrix:

| # | shift | Matrix | | | | | | | | | | | | |
|---|---|---|---|---|---|---|---|---|---|---|---|---|---|---|
| 1 | −24 | 58 | 0 | 0 | 0 | 0 | 0 | 0 | 0 | 0 | 0 | 0 | 0 | 0 |
| 2 | −23 | 57 | 0 | 0 | 0 | 0 | 0 | 0 | 0 | 0 | 0 | 0 | 0 | 0 |
| 3 | −21 | 0 | 58 | 0 | 0 | 0 | 0 | 0 | 0 | 0 | 0 | 0 | 0 | 0 |
| 4 | −20 | 0 | 57 | 58 | 0 | 0 | 0 | 0 | 0 | 0 | 0 | 0 | 0 | 0 |
| 5 | −19 | 0 | 0 | 57 | 0 | 0 | 0 | 0 | 0 | 0 | 0 | 0 | 0 | 0 |
| 6 | −18 | 52 | 0 | 0 | 0 | 0 | 0 | 0 | 0 | 0 | 0 | 0 | 0 | 0 |
| 7 | −16 | 50 | 0 | 0 | 0 | 0 | 0 | 0 | 0 | 0 | 0 | 0 | 0 | 0 |
| 8 | −15 | 0 | 52 | 0 | 58 | 0 | 0 | 0 | 0 | 0 | 0 | 0 | 0 | 0 |
| 9 | −14 | 48 | 0 | 52 | 57 | 0 | 0 | 0 | 0 | 0 | 0 | 0 | 0 | 0 |
| 10 | −13 | 47 | 50 | 0 | 0 | 58 | 0 | 0 | 0 | 0 | 0 | 0 | 0 | 0 |
| 11 | −12 | 46 | 0 | 50 | 0 | 57 | 0 | 0 | 0 | 0 | 0 | 0 | 0 | 0 |
| 12 | −11 | 45 | 48 | 0 | 0 | 0 | 58 | 0 | 0 | 0 | 0 | 0 | 0 | 0 |
| 13 | −10 | 0 | 47 | 48 | 0 | 0 | 57 | 58 | 0 | 0 | 0 | 0 | 0 | 0 |
| 14 | −9 | 0 | 46 | 47 | 52 | 0 | 0 | 57 | 58 | 0 | 0 | 0 | 0 | 0 |
| 15 | −8 | 42 | 45 | 46 | 0 | 0 | 0 | 0 | 57 | 58 | 0 | 0 | 0 | 0 |
| 16 | −7 | 41 | 0 | 45 | 50 | 52 | 0 | 0 | 0 | 57 | 0 | 0 | 0 | 0 |
| 17 | −5 | 39 | 42 | 0 | 48 | 50 | 52 | 0 | 0 | 0 | 58 | 0 | 0 | 0 |
| 18 | −4 | 38 | 41 | 42 | 47 | 0 | 0 | 52 | 0 | 0 | 57 | 0 | 0 | 0 |
| 19 | −3 | 0 | 0 | 41 | 46 | 48 | 50 | 0 | 52 | 0 | 0 | 0 | 0 | 0 |
| 20 | −2 | 0 | 39 | 0 | 45 | 47 | 0 | 50 | 0 | 52 | 0 | 58 | 0 | 0 |
| 21 | −1 | 0 | 38 | 39 | 0 | 46 | 48 | 0 | 50 | 0 | 0 | 57 | 58 | 0 |
| 22 | 0 | 0 | 0 | 38 | 0 | 45 | 47 | 48 | 0 | 50 | 0 | 0 | 57 | 0 |
| 23 | 1 | 33 | 0 | 0 | 42 | 0 | 46 | 47 | 48 | 0 | 52 | 0 | 0 | 0 |
| 24 | 2 | 0 | 0 | 0 | 41 | 0 | 45 | 46 | 47 | 48 | 0 | 0 | 0 | 0 |
| 25 | 3 | 0 | 0 | 0 | 0 | 42 | 0 | 45 | 46 | 47 | 50 | 0 | 0 | 0 |
| 26 | 4 | 0 | 33 | 0 | 39 | 41 | 0 | 0 | 45 | 46 | 0 | 52 | 0 | 58 |
| 27 | 5 | 0 | 0 | 33 | 38 | 0 | 42 | 0 | 0 | 45 | 48 | 0 | 52 | 57 |
| 28 | 6 | 0 | 0 | 0 | 0 | 39 | 41 | 42 | 0 | 0 | 47 | 50 | 0 | 0 |
| 29 | 7 | 0 | 0 | 0 | 0 | 38 | 0 | 41 | 42 | 0 | 46 | 0 | 50 | 0 |
| 30 | 8 | 0 | 0 | 0 | 0 | 0 | 39 | 0 | 41 | 42 | 45 | 48 | 0 | 0 |
| 31 | 9 | 0 | 0 | 0 | 0 | 0 | 38 | 39 | 0 | 41 | 0 | 47 | 48 | 0 |
| 32 | 10 | 0 | 0 | 0 | 33 | 0 | 0 | 38 | 39 | 0 | 0 | 46 | 47 | 52 |
| 33 | 11 | 0 | 0 | 0 | 0 | 0 | 0 | 0 | 38 | 39 | 42 | 45 | 46 | 0 |
| 34 | 12 | 0 | 0 | 0 | 0 | 33 | 0 | 0 | 0 | 38 | 41 | 0 | 45 | 50 |
| 35 | 14 | 0 | 0 | 0 | 0 | 0 | 33 | 0 | 0 | 0 | 39 | 42 | 0 | 48 |
| 36 | 15 | 0 | 0 | 0 | 0 | 0 | 0 | 33 | 0 | 0 | 38 | 41 | 42 | 47 |
| 37 | 16 | 0 | 0 | 0 | 0 | 0 | 0 | 0 | 33 | 0 | 0 | 0 | 41 | 46 |
| 38 | 17 | 0 | 0 | 0 | 0 | 0 | 0 | 0 | 0 | 33 | 0 | 39 | 0 | 45 |
| 39 | 18 | 0 | 0 | 0 | 0 | 0 | 0 | 0 | 0 | 0 | 0 | 38 | 39 | 0 |
| 40 | 19 | 0 | 0 | 0 | 0 | 0 | 0 | 0 | 0 | 0 | 0 | 0 | 38 | 0 |
| 41 | 20 | 0 | 0 | 0 | 0 | 0 | 0 | 0 | 0 | 0 | 33 | 0 | 0 | 42 |
| 42 | 21 | 0 | 0 | 0 | 0 | 0 | 0 | 0 | 0 | 0 | 0 | 0 | 0 | 41 |
| 43 | 23 | 0 | 0 | 0 | 0 | 0 | 0 | 0 | 0 | 0 | 0 | 33 | 0 | 39 |
| 44 | 24 | 0 | 0 | 0 | 0 | 0 | 0 | 0 | 0 | 0 | 0 | 0 | 33 | 38 |
| 45 | 29 | 0 | 0 | 0 | 0 | 0 | 0 | 0 | 0 | 0 | 0 | 0 | 0 | 33 |

We set a number $k$ of different rows to be used and for each $k$ we consider all possible combinations. If the number of rows in the matrix is $m$, then we need to consider $\frac{1}{1}$.

There are 13 ones in the second row. So, a new matrix must contain all indices of 1s, i.e. 33, 38, 39, 41, 42, 45, 46, 47, 48, 50, 52, 57, 58. For example, the following matrix (rows 21, 23, 26 of the original $(45 \times 13)$-matrix) has all 13 numbers.

| | 1 | 2 | 3 | 4 | 5 | 6 | 7 | 8 | 9 | 10 | 11 | 12 | 13 |
|---|---|---|---|---|---|---|---|---|---|---|---|---|---|
| −1 | 0 | 38 | 39 | 0 | 46 | 48 | 0 | 50 | 0 | 0 | 57 | 58 | 0 |
| 1 | 33 | 0 | 0 | 42 | 0 | 46 | 47 | 48 | 0 | 52 | 0 | 0 | 0 |
| 4 | 0 | 33 | 0 | 39 | 41 | 0 | 0 | 45 | 46 | 0 | 52 | 0 | 58 |

The first column has the only non-zero element, 33. So, all other elements 33 (second column) should be set to zero. The same is true, for elements 39 (3rd and 4th columns), 46 (5th, 6th and 9th columns), 52 (10th and 11th) and 58 (12th and 13th columns).

| | 1 | 2 | 3 | 4 | 5 | 6 | 7 | 8 | 9 | 10 | 11 | 12 | 13 |
|---|---|---|---|---|---|---|---|---|---|---|---|---|---|
| −1 | 0 | 38 | 39 | 0 | ~~46~~ | 48 | 0 | 50 | 0 | 0 | 57 | 58 | 0 |
| 1 | 33 | 0 | 0 | 42 | 0 | ~~46~~ | 47 | 48 | 0 | 52 | 0 | 0 | 0 |
| 4 | 0 | ~~33~~ | 0 | ~~39~~ | 41 | 0 | 0 | 45 | 46 | 0 | ~~52~~ | 0 | ~~58~~ |

and the matrix becomes

| | 1 | 2 | 3 | 4 | 5 | 6 | 7 | 8 | 9 | 10 | 11 | 12 | 13 |
|---|---|---|---|---|---|---|---|---|---|---|---|---|---|
| −1 | 0 | 38 | 39 | 0 | 0 | 48 | 0 | 50 | 0 | 0 | 57 | 58 | 0 |
| 1 | 33 | 0 | 0 | 42 | 0 | 0 | 47 | 48 | 0 | 52 | 0 | 0 | 0 |
| 4 | 0 | 0 | 0 | 0 | 41 | 0 | 0 | 45 | 46 | 0 | 0 | 0 | 0 |

The last column has only zero elements, so the chosen rows of the original $(45 \times 13)$-matrix are wrong.

Now we choose rows 15, 17, 36 and write down another matrix:

| | 1 | 2 | 3 | 4 | 5 | 6 | 7 | 8 | 9 | 10 | 11 | 12 | 13 |
|---|---|---|---|---|---|---|---|---|---|---|---|---|---|
| −8 | 42 | 45 | 46 | 0 | 0 | 0 | 0 | 57 | 58 | 0 | 0 | 0 | 0 |
| −5 | 39 | 42 | 0 | 48 | 50 | 52 | 0 | 0 | 0 | 58 | 0 | 0 | 0 |
| 15 | 0 | 0 | 0 | 0 | 0 | 0 | 33 | 0 | 0 | 38 | 41 | 42 | 47 |

Elements 58 (9th and 10th columns), 42 (1st, 2nd and 12th columns).

| | 1 | 2 | 3 | 4 | 5 | 6 | 7 | 8 | 9 | 10 | 11 | 12 | 13 |
|---|---|---|---|---|---|---|---|---|---|---|---|---|---|
| −8 | ~~42~~ | 45 | 46 | 0 | 0 | 0 | 0 | 57 | 58 | 0 | 0 | 0 | 0 |
| −5 | 39 | ~~42~~ | 0 | 48 | 50 | 52 | 0 | 0 | 0 | ~~58~~ | 0 | 0 | 0 |
| 15 | 0 | 0 | 0 | 0 | 0 | 0 | 33 | 0 | 0 | 38 | 41 | 42 | 47 |

Or by substituting zeros we obtain

| | 1 | 2 | 3 | 4 | 5 | 6 | 7 | 8 | 9 | 10 | 11 | 12 | 13 |
|---|---|---|---|---|---|---|---|---|---|---|---|---|---|
| −8 | 0 | 45 | 46 | 0 | 0 | 0 | 0 | 57 | 58 | 0 | 0 | 0 | 0 |
| −5 | 39 | 0 | 0 | 48 | 50 | 52 | 0 | 0 | 0 | 0 | 0 | 0 | 0 |
| 15 | 0 | 0 | 0 | 0 | 0 | 0 | 33 | 0 | 0 | 38 | 41 | 42 | 47 |

Each column has only one non-zero element.

For a given number of rows there may be no, one or multiple solutions. For example, there is another solution.

| | 1 | 2 | 3 | 4 | 5 | 6 | 7 | 8 | 9 | 10 | 11 | 12 | 13 |
|---|---|---|---|---|---|---|---|---|---|---|---|---|---|
| −4 | 38 | 41 | 42 | 47 | 0 | 0 | 52 | 0 | 0 | 57 | 0 | 0 | 0 |
| −1 | 0 | 38 | 39 | 0 | 46 | 48 | 0 | 50 | 0 | 0 | 57 | 58 | 0 |
| 17 | 0 | 0 | 0 | 0 | 0 | 0 | 0 | 0 | 33 | 0 | 39 | 0 | 45 |

The first step is processing for elements 38 and 57.

| | 1 | 2 | 3 | 4 | 5 | 6 | 7 | 8 | 9 | 10 | 11 | 12 | 13 |
|---|---|---|---|---|---|---|---|---|---|---|---|---|---|
| −4 | 38 | 41 | 42 | 47 | 0 | 0 | 52 | 0 | 0 | 57 | 0 | 0 | 0 |
| −1 | 0 | ~~38~~ | 39 | 0 | 46 | 48 | 0 | 50 | 0 | 0 | ~~57~~ | 58 | 0 |
| 17 | 0 | 0 | 0 | 0 | 0 | 0 | 0 | 0 | 33 | 0 | 39 | 0 | 45 |

| | 1 | 2 | 3 | 4 | 5 | 6 | 7 | 8 | 9 | 10 | 11 | 12 | 13 |
|---|---|---|---|---|---|---|---|---|---|---|---|---|---|
| −4 | 38 | 41 | 42 | 47 | 0 | 0 | 52 | 0 | 0 | 57 | 0 | 0 | 0 |
| −1 | 0 | 0 | 39 | 0 | 46 | 48 | 0 | 50 | 0 | 0 | 0 | 58 | 0 |
| 17 | 0 | 0 | 0 | 0 | 0 | 0 | 0 | 0 | 33 | 0 | 39 | 0 | 45 |

Now, we may process elements 39.

| | 1 | 2 | 3 | 4 | 5 | 6 | 7 | 8 | 9 | 10 | 11 | 12 | 13 |
|---|---|---|---|---|---|---|---|---|---|---|---|---|---|
| −4 | 38 | 41 | 42 | 47 | 0 | 0 | 52 | 0 | 0 | 57 | 0 | 0 | 0 |
| −1 | 0 | 0 | ~~39~~ | 0 | 46 | 48 | 0 | 50 | 0 | 0 | 0 | 58 | 0 |
| 17 | 0 | 0 | 0 | 0 | 0 | 0 | 0 | 0 | 33 | 0 | 39 | 0 | 45 |

| | 1 | 2 | 3 | 4 | 5 | 6 | 7 | 8 | 9 | 10 | 11 | 12 | 13 |
|---|---|---|---|---|---|---|---|---|---|---|---|---|---|
| −4 | 38 | 41 | 42 | 47 | 0 | 0 | 52 | 0 | 0 | 57 | 0 | 0 | 0 |
| −1 | 0 | 0 | 0 | 0 | 46 | 48 | 0 | 50 | 0 | 0 | 0 | 58 | 0 |
| 17 | 0 | 0 | 0 | 0 | 0 | 0 | 0 | 0 | 33 | 0 | 39 | 0 | 45 |

SIMD instructions can be written as

```
__m128i c, t, s;
c = _mm_set1_epi32(0xbcd8579b);
res[0] = _mm_and_si128(m[0], c);
c = _mm_set1_epi32(0x06006000);
t = _mm_and_si128(m[1], c);
s = _mm_srli_epi32(t, 8);
res[0] = _mm_or_si128(res[0], s);
c = _mm_set1_epi32(0x00150080);
t = _mm_and_si128(m[1], c);
s = _mm_srli_epi32(t, 5);
res[0] = _mm_or_si128(res[0], s);
c = _mm_set1_epi32(0x00008642);
t = _mm_and_si128(m[1], c);
s = _mm_slli_epi32(t, 15);
res[0] = _mm_or_si128(res[0], s);
```

for rows 15, 17, 36; and

```
__m128i c, t, s;
c = _mm_set1_epi32(0xbcd8579b);
res[0] = _mm_and_si128(m[0], c);
c = _mm_set1_epi32(0x02108640);
t = _mm_and_si128(m[1], c);
s = _mm_srli_epi32(t, 4);
res[0] = _mm_or_si128(res[0], s);
c = _mm_set1_epi32(0x04054000);
t = _mm_and_si128(m[1], c);
s = _mm_srli_epi32(t, 1);
res[0] = _mm_or_si128(res[0], s);
c = _mm_set1_epi32(0x00002082);
t = _mm_and_si128(m[1], c);
s = _mm_slli_epi32(t, 17);
res[0] = _mm_or_si128(res[0], s);
```

for rows 18, 21, 38.

Similar approaches can be used for spaced seeds of different lengths and weights.

#### S5. Read lengths as functions of seeds' period and weight

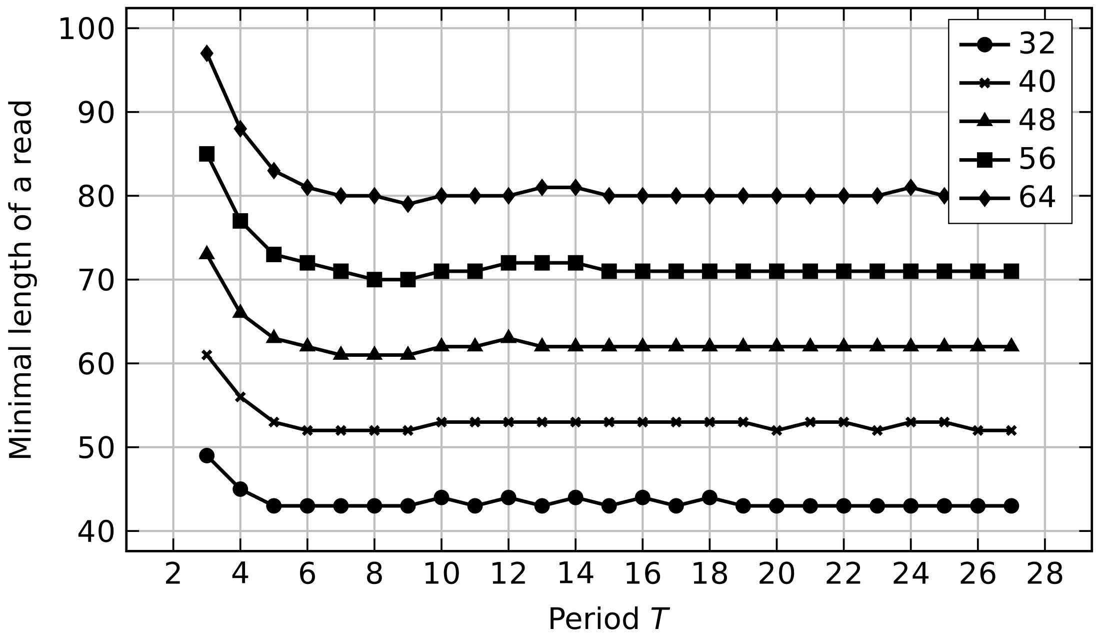


Number of mismatches: 1

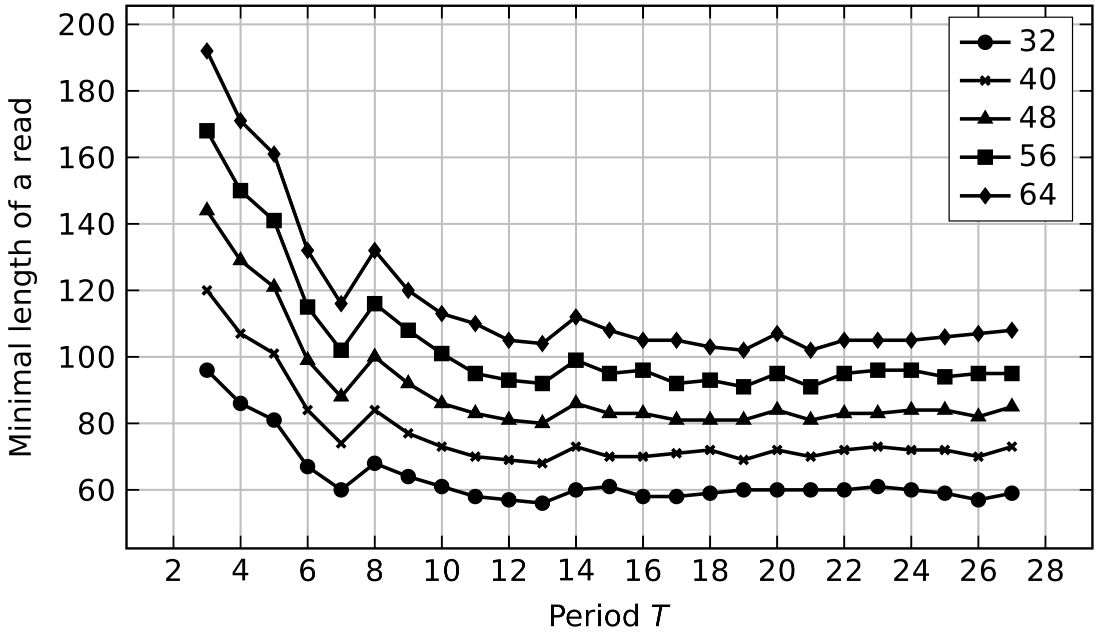


Number of mismatches: 2

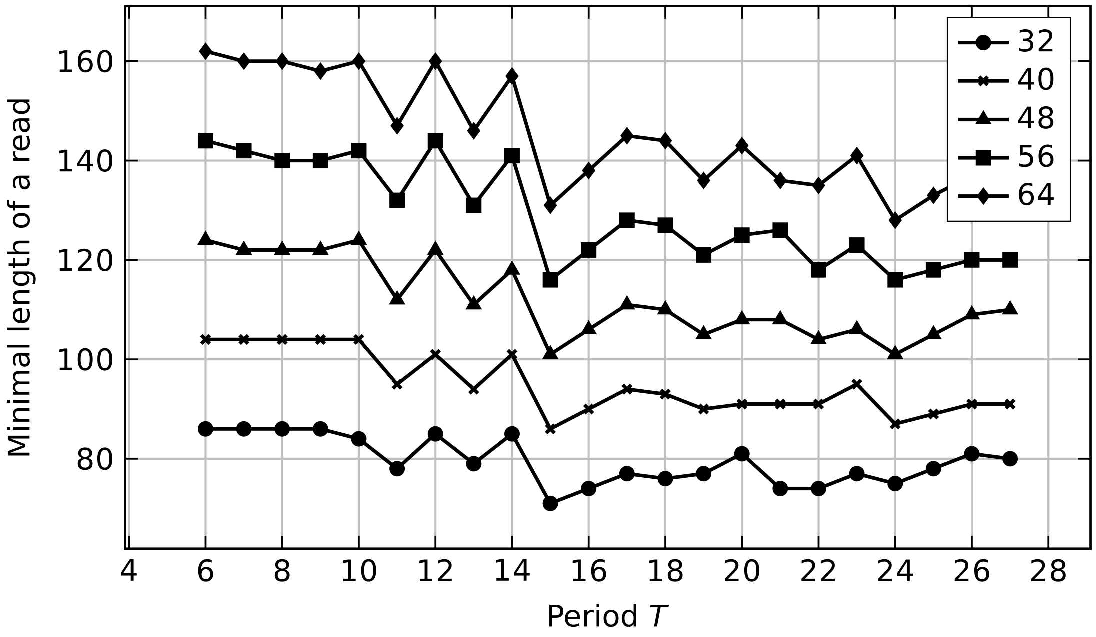

Number of mismatches: 3

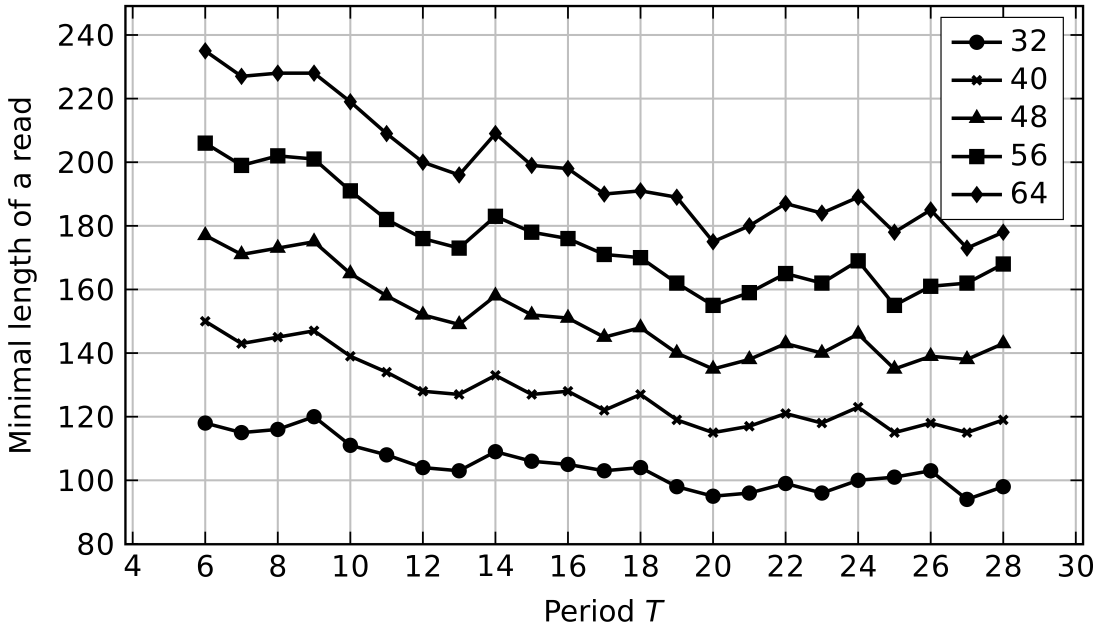

Number of mismatches: 4

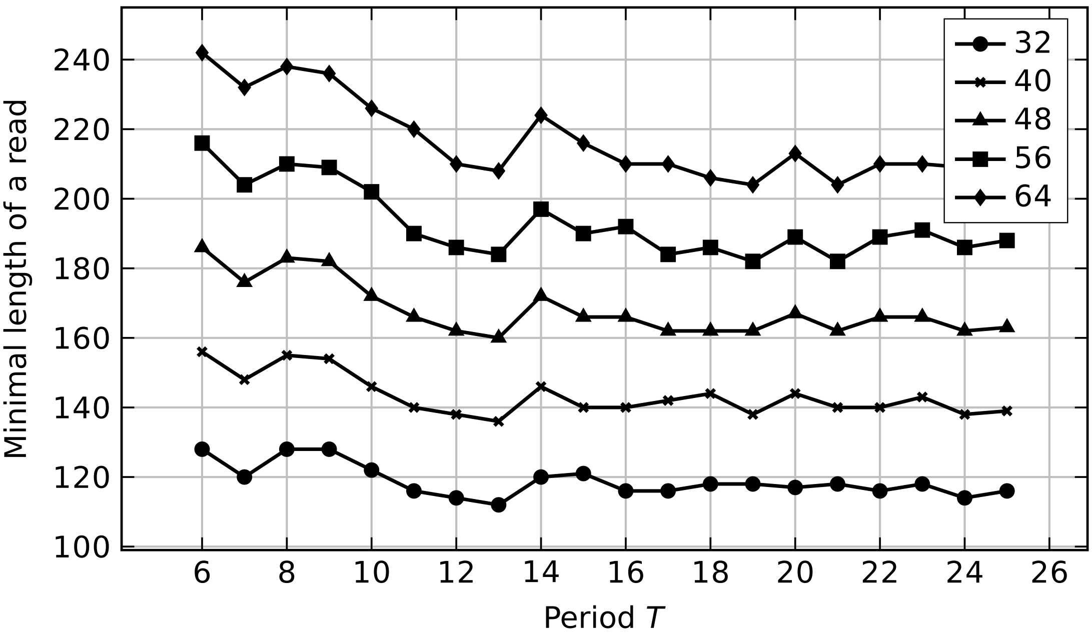


Number of mismatches: 5

### S6. Best spaced seeds

Spaced seeds of weight $w$ for reads permitting at most sub mismatches (the last column is the minimum length of a read required).

| Spaced seed | w | subs | len |
|---|---|---|---|
| 11110111101111011110111101111011110111101111 | 32 | 1 | 43 |
| 1111101110010111110111001011111011100101111 | 32 | 2 | 56 |
| 1111010110010001111010110010001111010110010001111010110 01 | 32 | 3 | 71 |
| 1110111001011100000010000000111011100101110000001000000011101110010111 | 32 | 4 | 94 |
| 1111101110010111110111001011111011100101111 | 32 | 5 | 112 |
| 111101011001000111101011001000111101011001000111101011001 | 32 | 6 | 135 |
| 111101011001000111101011001000111101011001000111101011001 | 32 | 7 | 142 |
| 1111101110010111110111001011111011100101111 | 32 | 8 | 168 |
| 1111101111101111101111101111101111101111101111 1 | 40 | 1 | 52 |
| 1111110101100111111010110011111101011001111110101100111 1 | 40 | 2 | 68 |
| 11110101100100011110101100100011110101100100011110101100100011110101 1001 | 40 | 3 | 86 |
| 1101111110000010100010001000011011111100000101000100010000110111111000001010001000100000 11011111 | 40 | 4 | 115 |
| 1111110101100111111010110011111101011001111110101100111 1 | 40 | 5 | 136 |
| 11110101100100011110101100100011110101100100011110101100100011110101 1001 | 40 | 6 | 165 |
| 11110101100100011110101100100011110101100100011110101100100011110101 1001 | 40 | 7 | 172 |
| 1111110111111011111101111110111111011111101111110111111 | 48 | 1 | 61 |
| 1110010111110111001011111011100101111101110010111110111001011111 0111 | 48 | 2 | 80 |
| 1111111000110100110010101111111000110100110010101111111000110100110010 10111111 | 48 | 3 | 101 |
| 1001011000110101000010010110001101010000100101100011010100001001011000110101000010010110001101<br>0100001001011000110101 | 48 | 4 | 135 |
| 1110010111110111001011111011100101111101110010111110111001011111 0111 | 48 | 5 | 160 |
| 1111111000110100110010101111111000110100110010101111111000110100110010 10111111 | 48 | 6 | 194 |
| 1111111000110100110010101111111000110100110010101111111000110100110010 10111111 | 48 | 7 | 202 |
| 111111101111111011111110111111101111111011111110111111101111111 | 56 | 1 | 70 |
| 11111111101101110001111111110110111000111111111011011100011111111101 10111 | 56 | 2 | 91 |
| 1110111110100111010010001110111110100111010010001110111110100111010010001110111110100111 01001 | 56 | 3 | 116 |
| 1000110010100111000010001100101001110000100011001010011100001000110010100111000010001100101001<br>110000100011001010011100001000110010100111 | 56 | 4 | 155 |
| 11111111101101110001111111110110111000111111111011011100011111111101 10111 | 56 | 5 | 182 |
| 1011100101111101110001001011100101111101110001001011100101111101110001001011100101111101110001 | 56 | 6 | 220 |
| 1111111101111111101111111101111111101111111101111111101111111101111111 1 | 64 | 1 | 79 |
| 111101011111111110110011110101111111111011001111010111111111101100111101011111111 11011 | 64 | 2 | 102 |
| 1110111110100111010010001110111110100111010010001110111110100111010010001110111110100111010010<br>00111011111 | 64 | 3 | 128 |
| 1101011110110001000000100000011010111101100010000001000000110101111011000100000010000001101011110110<br>00100000010000001101011110110001000000100000011010111101 1 | 64 | 4 | 173 |
| 111101011111111110110011110101111111111011001111010111111111101100111101011111111 11011 | 64 | 5 | 204 |

Minimum length of reads required for each seed for a given number of mismatches. Numbers in circles show the minimum values.

| Spaced seed (weight 32) | 1 | 2 | 3 | 4 | 5 | 6 | 7 | 8 |
|---|---|---|---|---|---|---|---|---|
| 1111110101100111111010110011111101011001111 1 | 50 | 56 | 99 | 106 | 112 | 153 | 160 | 168 |
| 1111101110010111110111001011111011100101111 | 49 | 56 | 98 | 103 | 112 | 151 | 157 | 168 |
| 1111110011010111111001101011111100110101111 1 | 50 | 56 | 98 | 105 | 112 | 151 | 158 | 168 |
| 111101011001000111101011001000111101011001000111101011001 | 61 | 67 | 71 | 123 | 127 | 135 | 142 | 183 |
| 1110111001011100000010000000111011100101110000001000000011101110010111 | 71 | 78 | 88 | 94 | 154 | 161 | 168 | 178 |

| Spaced seed (weight 40) | 1 | 2 | 3 | 4 | 5 | 6 | 7 |
|---|---|---|---|---|---|---|---|
| 11111101011001111110101100111111010110011111101011001111 | 62 | 68 | 120 | 128 | 136 | 182 | 191 |
| 11111011100101111101110010111110111001011111011100101111 | 61 | 68 | 121 | 127 | 136 | 184 | 193 |
| 11111100110101111110011010111111001101011111100110101111 | 62 | 68 | 122 | 130 | 136 | 187 | 193 |
| 11111010011101111101001110111110100111011111010011101111 | 61 | 68 | 120 | 128 | 136 | 184 | 192 |
| 111101011001000111101011001000111101011001000111101011001000111101011001 | 76 | 82 | 86 | 153 | 157 | 165 | 172 |
| 100011001010011100001000110010100111000010001100101001110000100011001010011100001000110010100111 | 99 | 102 | 110 | 115 | 201 | 207 | 213 |
| 100101100011010100001001011000110101000010010110001101010000100101100011010100001001011000110101 | 98 | 103 | 110 | 115 | 201 | 206 | 213 |
| 100111001010001100001001110010100011000010011100101000110000100111001010001100001001110010100011 | 99 | 103 | 109 | 115 | 201 | 207 | 213 |
| 110101100010010100001101011000100101000011010110001001010000110101100010010100001101011000100101 | 98 | 104 | 108 | 115 | 202 | 209 | 214 |
| 1010111001001111000000000010101110010011110000000000101011100100111100000000001010111001001111 | 95 | 99 | 111 | 115 | 196 | 205 | 210 |
| 1011100111001011000000000010111001110010110000000000101110011100101100000000001011100111001011 | 94 | 102 | 112 | 115 | 198 | 204 | 210 |
| 1011101101100011000000000010111011011000110000000000101110110110001100000000001011101101100011 | 94 | 103 | 110 | 115 | 197 | 204 | 209 |
| 10111111000000101000110000000101111110000001010001100000001011111100000010100011000000010111111 | 95 | 101 | 109 | 115 | 200 | 206 | 212 |
| 11011111100001010001000100000110111111000010100010001000001101111110000101000100010000011011111 | 94 | 99 | 108 | 115 | 197 | 203 | 211 |
| 11011111100010010000010100000011011111100010010000010100000011011111100010010000010100000011011111 | 94 | 99 | 108 | 115 | 197 | 205 | 213 |
| 11011111100011001000001000000011011111100011001000001000000011011111100011001000001000000011011111 | 94 | 99 | 110 | 115 | 197 | 205 | 211 |
| 11101111100000001001100000001011101111100000001001100000001011101111100000001001100000001011101111 | 93 | 100 | 108 | 115 | 196 | 202 | 210 |
| 11101111010000001100001000000111011110100000011000010000001110111101000000110000100000011101111 | 93 | 100 | 108 | 115 | 196 | 203 | 210 |

| Spaced seed (weight 48) | 1 | 2 | 3 | 4 | 5 | 6 | 7 |
|---|---|---|---|---|---|---|---|
| 11100101111101110010111110111001011111011100101111101110010111110111 | 73 | 80 | 142 | 149 | 160 | 215 | 227 |
| 11111010011101111101001110111110100111011111010011101111101001110111 | 73 | 80 | 144 | 153 | 160 | 217 | 229 |
| 11111011100101111101110010111110111001011111011100101111101110010111 | 73 | 80 | 142 | 150 | 160 | 218 | 228 |
| 11111100110101111110011010111111001101011111100110101111110011010111 | 74 | 80 | 144 | 153 | 160 | 218 | 228 |
| 11111101011001111110101100111111010110011111101011001111110101100111 | 74 | 80 | 144 | 153 | 160 | 217 | 228 |
| 100110101111000100110101111000100110101111000100110101111000100110101111000100110101111 | 91 | 97 | 101 | 183 | 187 | 195 | 202 |
| 111111101010011001011000111111101010011001011000111111101010011001011000111111 | 85 | 94 | 101 | 176 | 184 | 195 | 202 |
| 11111110001101001100101011111111000110100110010101111111100011010011001010111111 | 85 | 94 | 101 | 176 | 184 | 194 | 202 |
| 10001100101001110000100011001010011100001000110010100111000010001100101001110000100011001010011100001000110010100111 | 119 | 122 | 130 | 135 | 241 | 248 | 253 |
| 10010110001101010000100101100011010100001001011000110101000010010110001101010000100101100011010100001001011000110101 | 118 | 123 | 130 | 135 | 241 | 246 | 253 |
| 10011100101000110000100111001010001100001001110010100011000010011100101000110000100111001010001100001001110010100011 | 119 | 123 | 129 | 135 | 241 | 247 | 254 |
| 10100100011010110000101001000110101100001010010001101011000010100100011010110000101001000110101100001010010001101011 | 118 | 124 | 128 | 135 | 243 | 249 | 254 |
| 1011101101100011000000000010111011011000110000000000101110110110001100000000001011101101100011000000000010111011011 | 114 | 123 | 130 | 135 | 240 | 248 | 255 |

| Spaced seed (weight 56) | 1 | 2 | 3 | 4 | 5 | 6 | 7 |
|---|---|---|---|---|---|---|---|
| 1111111110110111000111111111011011100011111111101101110001111111110110111 | 82 | 91 | 157 | 170 | 182 | 240 | 254 |
| 11111111101100111101011111111101100111101011111111101100111101011111111 | 80 | 91 | 158 | 168 | 182 | 244 | 253 |
| 11111111101011110011011111111101011110011011111111101011110011011111111 | 80 | 91 | 159 | 171 | 182 | 243 | 255 |
| 111101011001000111101011001000111101011001000111101011001000111101011001000111101011001000111101011001 | 106 | 112 | 116 | 213 | 217 | 225 | 232 |
| 111011111010011101001000111011111010011101001000111011111010011101001000111011111010011101001 | 98 | 109 | 116 | 199 | 210 | 221 | 232 |
| 111111101010011001011000111111101010011001011000111111101010011001011000111111101010011001011 | 100 | 109 | 116 | 200 | 211 | 223 | 232 |
| 1000110010100111000010001100101001110000100011001010011100001000110010100111000010001100101001110000100011001010011100001000110010100111 | 139 | 142 | 150 | 155 | 279 | 287 | 292 |
| 1001011000110101000010010110001101010000100101100011010100001001011000110101000010010110001101010000100101100011010100001001011000110101 | 138 | 143 | 150 | 155 | 280 | 285 | 291 |
| 1001110010100011000010011100101000110000100111001010001100001001110010100011000010011100101000110000100111001010001100001001110010100011 | 139 | 143 | 149 | 155 | 280 | 287 | 293 |
| 1010010001101011000010100100011010110000101001000110101100001010010001101011000010100100011010110000101001000110101100001010010001101011 | 138 | 144 | 148 | 155 | 282 | 286 | 294 |
| 1111110010100010000100000011111100101000100001000000111111001010001000010000001111110010100010000100000011111100101000100001000000111111 | 137 | 143 | 149 | 155 | 285 | 290 | 297 |
| 1011100101111101110001001011100101111101110001001011100101111101110001001011100101111101110001 | 99 | 110 | 117 | 199 | 209 | 220 | 234 |

| Spaced seed (weight 64) | 1 | 2 | 3 | 4 | 5 | 6 | 7 |
|---|---|---|---|---|---|---|---|
| 1111010111111111011001111010111111111011001111010111111111011001111010111111111011 | 91 | 102 | 176 | 188 | 204 | 268 | 284 |
| 111111110101110011011111111010111001101111111101011100110111111110101110011011111111 | 92 | 102 | 184 | 193 | 204 | 279 | 290 |
| 111111111000111011011111111100011101101111111110001110110111111111000111011011111111 | 93 | 102 | 184 | 194 | 204 | 280 | 290 |
| 111111111001101101011111111100110110101111111110011011010111111111001101101011111111 | 93 | 102 | 184 | 194 | 204 | 280 | 291 |
| 111111111010110110011111111101011011001111111110101101100111111111010110110011111111 | 93 | 102 | 185 | 195 | 204 | 282 | 290 |
| 111111111011011100011111111101101110001111111110110111000111111111011011100011111111 | 93 | 102 | 185 | 195 | 204 | 280 | 292 |
| 111011111010011101001000111011111010011101001000111011111010011101001000111011111010 011101001000111011111 | 110 | 121 | 128 | 225 | 234 | 247 | 256 |
| 111111101010011001011000111111101010011001011000111111101010011001011000111111101010 0110010110001111111101 | 112 | 121 | 128 | 229 | 238 | 247 | 256 |
| 111110101011001100000000000111110101011001100000000000111110101011001100000000000111 110101011001100000000000111110101011001100000000000111110101011 | 152 | 160 | 167 | 173 | 313 | 319 | 325 |
| 110101111011000100000100000110101111011000100000100000110101111011000100000100000110 101111011000100000100000110101111011000100000100000110101111011 | 151 | 160 | 167 | 173 | 312 | 318 | 326 |

### S7. SIMD instructions

#### S7..1 Weight 32

- 111101111011110111101111011110111101111

  **Input seed:**

  ```
  AAAA0AAA A0AAAA0A AAA0AAAA 0AAAA0AA
  ED0CBED0 00000000 00000000 00000000
  ```

  **Output seed:**

  ```
  AAAABAAA ACAAAADA AAADAAAA EAAAAEAA

  Row  1: AAAA0AAAA0AAAA0AAAA0AAAA0AAAA0AA

  Ext  B: 0000B000000000000000000000000000
  Ext  C:       000C0000000000000000000000000000
  Ext  D:              0D0000D0000000000000000000000000
  Ext  E:                         E0000E00000000000000000000000000

  __m128i c, t, s;
  c = _mm_set1_epi32(0xdef7bdef);
  res[0] = _mm_and_si128(m[0], c);
  c = _mm_set1_epi32(0x00000010);
  t = _mm_and_si128(m[1], c);
  res[0] = _mm_or_si128(res[0], t);
  c = _mm_set1_epi32(0x00000008);
  t = _mm_and_si128(m[1], c);
  s = _mm_slli_epi32(t, 6);
  res[0] = _mm_or_si128(res[0], s);
  c = _mm_set1_epi32(0x00000042);
  t = _mm_and_si128(m[1], c);
  s = _mm_slli_epi32(t, 13);
  res[0] = _mm_or_si128(res[0], s);
  c = _mm_set1_epi32(0x00000021);
  t = _mm_and_si128(m[1], c);
  s = _mm_slli_epi32(t, 24);
  res[0] = _mm_or_si128(res[0], s);
  ```

- 11111011100101111101110010111110111001011111

  **Input seed:**

  ```
  AAAAA0AA A00A0AAA AA0AAA00 A0AAAAA0
  DCD00B0C DBBC0000 00000000 00000000
  ```

  **Output seed:**

  ```
  AAAAABAA ABBACAAA AACAAACD ADAAAAAD

  Row  1: AAAAA0AAA00A0AAAAA0AAA00A0AAAAA0

  Ext  B: 00000B000BB000000000000000000000
  Ext  C:            0C00000C000C00000000000000000000
  Ext  D:                        0D0D00000D0000000000000000000000

  __m128i c, t, s;
  c = _mm_set1_epi32(0x7d3be9df);
  res[0] = _mm_and_si128(m[0], c);
  c = _mm_set1_epi32(0x00000620);
  ```

```
t = _mm_and_si128(m[1], c);
res[0] = _mm_or_si128(res[0], t);
c = _mm_set1_epi32(0x00000882);
t = _mm_and_si128(m[1], c);
s = _mm_slli_epi32(t, 11);
res[0] = _mm_or_si128(res[0], s);
c = _mm_set1_epi32(0x00000105);
t = _mm_and_si128(m[1], c);
s = _mm_slli_epi32(t, 23);
res[0] = _mm_or_si128(res[0], s);
```

- 111101011001000111101011001000111101011001000111101011001

  **Input seed:**

  ```
  AAAA0A0A A00A000A AAA0A0AA 00A000AA
  ED0D0EC0 0C000CED D0D0DC00 B0000000
  ```

  **Output seed:**

  ```
  AAAABACA ACDADCEA AAAEACAA DDADEDAA
  ```

  ```
  Row  1:                     AAAA0A0AA00A000AAAA0A0AA00A000AA

  Ext  B: 000000000000000000000000B0000000
  Ext  C:                     000000C00C000C0000000C0000000000
  Ext  D:                             0D0D00000000000DD0D0D00000000000
  Ext  E:                                  E0000E00000000E00000000000000000
  ```

  ```
  __m128i c, t, s;
  c = _mm_set1_epi32(0xc4d789af);
  res[0] = _mm_and_si128(m[0], c);
  c = _mm_set1_epi32(0x01000000);
  t = _mm_and_si128(m[1], c);
  s = _mm_srli_epi32(t, 20);
  res[0] = _mm_or_si128(res[0], s);
  c = _mm_set1_epi32(0x00202240);
  t = _mm_and_si128(m[1], c);
  res[0] = _mm_or_si128(res[0], t);
  c = _mm_set1_epi32(0x0015800a);
  t = _mm_and_si128(m[1], c);
  s = _mm_slli_epi32(t, 9);
  res[0] = _mm_or_si128(res[0], s);
  c = _mm_set1_epi32(0x00004021);
  t = _mm_and_si128(m[1], c);
  s = _mm_slli_epi32(t, 14);
  res[0] = _mm_or_si128(res[0], s);
  ```

- 11101110010111000000100000000111011100101110000001000000001110111001011 1

  **Input seed:**

  ```
  AAA0AAA0 0A0AAA00 000A0000 000AAA0A
  DE00D0ED C00000D0 000000CB C0BDB00B
  0FFF0000 00000000 00000000 00000000
  ```

  **Output seed:**

  ```
  AAADAAAD CADAAAFF FDBAEBCB CEBAAADA
  ```

```
Row  1:      AAA0AAA00A0AAA00000A0000000AAA0A

Ext  B: 00000000000000000000000B00B0B00B
Ext  C:      00000000C0000000000000C0C0000000
Ext  D:         D000D00D000000D000000000000D0000
Ext  E:                         0E0000E0000000000000000000000000
Ext  F:                   0FFF0000000000000000000000000000

__m128i c, t, s;
c = _mm_set1_epi32(0xb8083a77);
res[0] = _mm_and_si128(m[0], c);
c = _mm_set1_epi32(0x94800000);
t = _mm_and_si128(m[1], c);
s = _mm_srli_epi32(t, 5);
res[0] = _mm_or_si128(res[0], s);
c = _mm_set1_epi32(0x01400100);
t = _mm_and_si128(m[1], c);
res[0] = _mm_or_si128(res[0], t);
c = _mm_set1_epi32(0x08004091);
t = _mm_and_si128(m[1], c);
s = _mm_slli_epi32(t, 3);
res[0] = _mm_or_si128(res[0], s);
c = _mm_set1_epi32(0x00000042);
t = _mm_and_si128(m[1], c);
s = _mm_slli_epi32(t, 19);
res[0] = _mm_or_si128(res[0], s);
c = _mm_set1_epi32(0x0000000e);
t = _mm_and_si128(m[2], c);
s = _mm_slli_epi32(t, 13);
res[0] = _mm_or_si128(res[0], s);
```

#### S7..2 Weight 40

- 11111011111011111011111011111011111011111011111
  **Input seed:**

```
AAAAA0AA AAA0AAAA A0AAAAA0 AAAAA0AA
BBB0BBBB F0GDECF0 00000000 00000000
```

**Output seed:**

```
AAAAACAA AAADAAAA AEAAAAAF AAAAAFAA
BBBGBBBB 00000000 00000000 00000000
```

```
Row  1:         AAAAA0AAAAA0AAAAA0AAAAA0AAAAA0AA

Ext  C: 0000000000000C000000000000000000
Ext  D:         00000000000D00000000000000000000
Ext  E:              000000000000E0000000000000000000
Ext  F:                        00000000F00000F00000000000000000

Row  2:         BBB0BBBB000000000000000000000000

Ext  G:  0000000000G000000000000000000000
```

```
__m128i c, t, s;
c = _mm_set1_epi32(0xdf7df7df);
res[0] = _mm_and_si128(m[0], c);
c = _mm_set1_epi32(0x00002000);
t = _mm_and_si128(m[1], c);
s = _mm_srli_epi32(t, 8);
res[0] = _mm_or_si128(res[0], s);
c = _mm_set1_epi32(0x00000800);
t = _mm_and_si128(m[1], c);
res[0] = _mm_or_si128(res[0], t);
c = _mm_set1_epi32(0x00001000);
t = _mm_and_si128(m[1], c);
s = _mm_slli_epi32(t, 5);
res[0] = _mm_or_si128(res[0], s);
c = _mm_set1_epi32(0x00004100);
t = _mm_and_si128(m[1], c);
s = _mm_slli_epi32(t, 15);
res[0] = _mm_or_si128(res[0], s);

c = _mm_set1_epi32(0x000000f7);
res[1] = _mm_and_si128(m[1], c);
c = _mm_set1_epi32(0x00000400);
t = _mm_and_si128(m[1], c);
s = _mm_srli_epi32(t, 7);
res[1] = _mm_or_si128(res[1], s);
```

- 111111010110011111101011001111110101100111111010110011111
  **Input seed:**

```
AAAAAA0A 0AA00AAA AAA0A0AA 00AAAAAA
0B0BB00B DCFCF0C0 FD00DDCE 00000000
```

**Output seed:**

```
AAAAAACA CAACDAAA AAACADAA DDAAAAAA
FBFBBEFB 00000000 00000000 00000000
```

```
Row  1:                   AAAAAA0A0AA00AAAAAA0A0AA00AAAAAA

Ext  C:                000000000C0C00C0000000C000000000
Ext  D:                       00000000D00000000D00DD0000000000

Row  2:                   0B0BB00B000000000000000000000000

Ext  E: 00000000000000000000000E00000000
Ext  F:         0000000000F0F000F000000000000000
```

```
__m128i c, t, s;
c = _mm_set1_epi32(0xfcd7e6bf);
res[0] = _mm_and_si128(m[0], c);
c = _mm_set1_epi32(0x00404a00);
t = _mm_and_si128(m[1], c);
s = _mm_srli_epi32(t, 3);
res[0] = _mm_or_si128(res[0], s);
c = _mm_set1_epi32(0x00320100);
t = _mm_and_si128(m[1], c);
s = _mm_slli_epi32(t, 4);
res[0] = _mm_or_si128(res[0], s);

c = _mm_set1_epi32(0x0000009a);
res[1] = _mm_and_si128(m[1], c);
c = _mm_set1_epi32(0x00800000);
t = _mm_and_si128(m[1], c);
s = _mm_srli_epi32(t, 18);
res[1] = _mm_or_si128(res[1], s);
c = _mm_set1_epi32(0x00011400);
t = _mm_and_si128(m[1], c);
s = _mm_srli_epi32(t, 10);
res[1] = _mm_or_si128(res[1], s);
```

- 111101011001000111101011001000111101011001000111101011001000111101011001

  **Input seed:**

```
AAAA0A0A A00A000A AAA0A0AA 00A000AA
BB0B0BB0 0E000ECG C0D0CE00 C000ECDD
0F0FH00H 00000000 00000000 00000000
```

  **Output seed:**

```
AAAACACA AECADECA AAACAEAA DDAFEFAA
BBGBHBBH 00000000 00000000 00000000
```

```
Row  1:              AAAA0A0AA00A000AAAA0A0AA00A000AA

Ext  C:    00000000000000C0C000C000C0000C00
Ext  D:        000000000000000000D00000000000DD
Ext  E:              000000000E000E0000000E000000E000
Ext  F:                                 0F0F0000000000000000000000000000

Row  2:              BB0B0BB0000000000000000000000000

Ext  G: 000000000000000G0000000000000000
Ext  H:              0000H00H000000000000000000000000
```

```
__m128i c, t, s;
c = _mm_set1_epi32(0xc4d789af);
res[0] = _mm_and_si128(m[0], c);
```

```
c = _mm_set1_epi32(0x21114000);
t = _mm_and_si128(m[1], c);
s = _mm_srli_epi32(t, 10);
res[0] = _mm_or_si128(res[0], s);
c = _mm_set1_epi32(0xc0040000);
t = _mm_and_si128(m[1], c);
s = _mm_srli_epi32(t, 6);
res[0] = _mm_or_si128(res[0], s);
c = _mm_set1_epi32(0x10202200);
t = _mm_and_si128(m[1], c);
res[0] = _mm_or_si128(res[0], t);
c = _mm_set1_epi32(0x0000000a);
t = _mm_and_si128(m[2], c);
s = _mm_slli_epi32(t, 26);
res[0] = _mm_or_si128(res[0], s);

c = _mm_set1_epi32(0x0000006b);
res[1] = _mm_and_si128(m[1], c);
c = _mm_set1_epi32(0x00008000);
t = _mm_and_si128(m[1], c);
s = _mm_srli_epi32(t, 13);
res[1] = _mm_or_si128(res[1], s);
c = _mm_set1_epi32(0x00000090);
t = _mm_and_si128(m[2], c);
res[1] = _mm_or_si128(res[1], t);
```

- 11011111000010100010001000001101111100001010001000100000110111110000101000100010000011011111

  **Input seed:**

```
AA0AAAAA 0000A0A0 00A000A0 000AA0AA
BBB0000B 0D000E00 0D0000DE 0CEDDE00
00G0G000 F000G000 0FG0HHHH F0000000
```

  **Output seed:**

```
AADAAAAA FGDGAEAD CFAGDDAE FGEAAEAA
BBBHHHHB 00000000 00000000 00000000

Row  1:                  AA0AAAAA0000A0A000A000A0000AA0AA

Ext  C:         0000000000000000000000000C000000
Ext  D:           000000000D0000000D0000D0000DD000
Ext  E:                  0000000000000E000000000E00E00E00
Ext  F:                  00000000F00000000F000000F0000000
Ext  G:                         00G0G0000000G00000G0000000000000

Row  2:                  BBB0000B000000000000000000000000

Ext  H: 00000000000000000000HHHH00000000

__m128i c, t, s;
c = _mm_set1_epi32(0xd84450fb);
res[0] = _mm_and_si128(m[0], c);
c = _mm_set1_epi32(0x02000000);
t = _mm_and_si128(m[1], c);
s = _mm_srli_epi32(t, 9);
res[0] = _mm_or_si128(res[0], s);
c = _mm_set1_epi32(0x18420200);
t = _mm_and_si128(m[1], c);
s = _mm_srli_epi32(t, 7);
```

```
res[0] = _mm_or_si128(res[0], s);
c = _mm_set1_epi32(0x24802000);
t = _mm_and_si128(m[1], c);
res[0] = _mm_or_si128(res[0], t);
c = _mm_set1_epi32(0x01020100);
t = _mm_and_si128(m[2], c);
res[0] = _mm_or_si128(res[0], t);
c = _mm_set1_epi32(0x00041014);
t = _mm_and_si128(m[2], c);
s = _mm_slli_epi32(t, 7);
res[0] = _mm_or_si128(res[0], s);

c = _mm_set1_epi32(0x00000087);
res[1] = _mm_and_si128(m[1], c);
c = _mm_set1_epi32(0x00f00000);
t = _mm_and_si128(m[2], c);
s = _mm_srli_epi32(t, 17);
res[1] = _mm_or_si128(res[1], s);
```

- 1111010110010001111010110010001111010110010001111010110010001111 01011001

**Input seed:**

```
AAAA0A0A A00A000A AAA0A0AA 00A000AA
BB0B0BB0 0E000DED D0C0CE00 E000DEDD
0H0HF00G 00000000 00000000 00000000
```

**Output seed:**

```
AAAACACA AEDADDEA AAAFAEAA EDADDEAA
BBHBHBBG 00000000 00000000 00000000

Row  1:               AAAA0A0AA00A000AAAA0A0AA00A000AA

Ext  C: 000000000000000000C0C00000000000
Ext  D:            0000000000000D0DD00000000000D0DD
Ext  E:               000000000E0000E000000E00E0000E00
Ext  F:                              0000F000000000000000000000000000

Row  2:               BB0B0BB0000000000000000000000000

Ext  G:               0000000G000000000000000000000000
Ext  H:                0H0H0000000000000000000000000000

__m128i c, t, s;
c = _mm_set1_epi32(0xc4d789af);
res[0] = _mm_and_si128(m[0], c);
c = _mm_set1_epi32(0x00140000);
t = _mm_and_si128(m[1], c);
s = _mm_srli_epi32(t, 14);
res[0] = _mm_or_si128(res[0], s);
c = _mm_set1_epi32(0xd001a000);
t = _mm_and_si128(m[1], c);
s = _mm_srli_epi32(t, 3);
res[0] = _mm_or_si128(res[0], s);
c = _mm_set1_epi32(0x21204200);
t = _mm_and_si128(m[1], c);
res[0] = _mm_or_si128(res[0], t);
c = _mm_set1_epi32(0x00000010);
t = _mm_and_si128(m[2], c);
s = _mm_slli_epi32(t, 15);
```

```
res[0] = _mm_or_si128(res[0], s);

c = _mm_set1_epi32(0x0000006b);
res[1] = _mm_and_si128(m[1], c);
c = _mm_set1_epi32(0x00000080);
t = _mm_and_si128(m[2], c);
res[1] = _mm_or_si128(res[1], t);
c = _mm_set1_epi32(0x0000000a);
t = _mm_and_si128(m[2], c);
s = _mm_slli_epi32(t, 1);
res[1] = _mm_or_si128(res[1], s);
```

#### S7..3 Weight 48

- 1111110111111011111101111110111111011111101111110111111

  **Input seed:**

```
AAAAAA0A AAAAA0AA AAAA0AAA AAA0AAAA
BB0BBBBB B0BBBBBB 0HGFEDC0 00000000
```

  **Output seed:**

```
AAAAAACA AAAAADAA AAAAEAAA AAAFAAAA
BBGBBBBB BHBBBBBB 00000000 00000000

Row  1:                 AAAAAA0AAAAAA0AAAAAA0AAAAAA0AAAA

Ext  C: 0000000000000000000000C000000000
Ext  D:         000000000000000000000D0000000000
Ext  E:                 00000000000000000000E00000000000
Ext  F:                         0000000000000000000F000000000000

Row  2:                 BB0BBBBBB0BBBBBB0000000000000000

Ext  G: 000000000000000000G0000000000000
Ext  H:         00000000000000000H00000000000000

__m128i c, t, s;
c = _mm_set1_epi32(0xf7efdfbf);
res[0] = _mm_and_si128(m[0], c);
c = _mm_set1_epi32(0x00400000);
t = _mm_and_si128(m[1], c);
s = _mm_srli_epi32(t, 16);
res[0] = _mm_or_si128(res[0], s);
c = _mm_set1_epi32(0x00200000);
t = _mm_and_si128(m[1], c);
s = _mm_srli_epi32(t, 8);
res[0] = _mm_or_si128(res[0], s);
c = _mm_set1_epi32(0x00100000);
t = _mm_and_si128(m[1], c);
res[0] = _mm_or_si128(res[0], t);
c = _mm_set1_epi32(0x00080000);
t = _mm_and_si128(m[1], c);
s = _mm_slli_epi32(t, 8);
res[0] = _mm_or_si128(res[0], s);

c = _mm_set1_epi32(0x0000fdfb);
res[1] = _mm_and_si128(m[1], c);
c = _mm_set1_epi32(0x00040000);
t = _mm_and_si128(m[1], c);
s = _mm_srli_epi32(t, 16);
res[1] = _mm_or_si128(res[1], s);
c = _mm_set1_epi32(0x00020000);
t = _mm_and_si128(m[1], c);
s = _mm_srli_epi32(t, 8);
res[1] = _mm_or_si128(res[1], s);
```

- 11100101111101110010111110111001011111011100101111101110010111110111

  **Input seed:**

```
AAA00A0A AAAA0AAA 00A0AAAA A0AAA00A
0BBBBB0B BB00B0BB EED0CGC0 0E0GCEEG
0HFH0000 00000000 00000000 00000000
```

**Output seed:**

```
AAADCACA AAAACAAA EEAFAAAA AEAAAEEA
GBBBBBGB BBGHBHBB 00000000 00000000

Row  1:                      AAA00A0AAAAA0AAA00A0AAAAA0AAA00A

Ext  C:      00000000000000000000C0C00000C000
Ext  D:       000000000000000000D0000000000000
Ext  E:                      0000000000000000EE0000000E000EE0
Ext  F:                                      00F00000000000000000000000000000

Row  2:                      0BBBBB0BBB00B0BB0000000000000000

Ext  G: 000000000000000000000G00000G000G
Ext  H:                                0H0H0000000000000000000000000000
```

```
__m128i c, t, s;
c = _mm_set1_epi32(0x9df4efa7);
res[0] = _mm_and_si128(m[0], c);
c = _mm_set1_epi32(0x10500000);
t = _mm_and_si128(m[1], c);
s = _mm_srli_epi32(t, 16);
res[0] = _mm_or_si128(res[0], s);
c = _mm_set1_epi32(0x00040000);
t = _mm_and_si128(m[1], c);
s = _mm_srli_epi32(t, 15);
res[0] = _mm_or_si128(res[0], s);
c = _mm_set1_epi32(0x62030000);
t = _mm_and_si128(m[1], c);
res[0] = _mm_or_si128(res[0], t);
c = _mm_set1_epi32(0x00000004);
t = _mm_and_si128(m[2], c);
s = _mm_slli_epi32(t, 17);
res[0] = _mm_or_si128(res[0], s);

c = _mm_set1_epi32(0x0000d3be);
res[1] = _mm_and_si128(m[1], c);
c = _mm_set1_epi32(0x88200000);
t = _mm_and_si128(m[1], c);
s = _mm_srli_epi32(t, 21);
res[1] = _mm_or_si128(res[1], s);
c = _mm_set1_epi32(0x0000000a);
t = _mm_and_si128(m[2], c);
s = _mm_slli_epi32(t, 10);
res[1] = _mm_or_si128(res[1], s);
```

- 1111111000110100110010101111111000110100110010101111111000110100110010101111111

  **Input seed:**

  ```
  AAAAAAA0 00AA0A00 AA00A0A0 AAAAAAA0
  00BB0B00 BB00B0B0 EDEGDDC0 00EG0C00
  HH00H0H0 FFHHFH00 00000000 00000000
  ```

  **Output seed:**

  ```
  AAAAAAAC FFAAFACD AADDAEAE AAAAAAAE
  HHBBHBHG BBHHBHBG 00000000 00000000
  ```

```
Row  1:                AAAAAAA000AA0A00AA00A0A0AAAAAAA0

Ext  C: 0000000000000000000000C000000C00
Ext  D:              00000000000000000D00DD0000000000
Ext  E:                     0000000000000000E0E0000000E00000
Ext  F:                00000000FF00F0000000000000000000

Row  2:                00BB0B00BB00B0B00000000000000000

Ext  G:    0000000000000000000G0000000G0000
Ext  H:                HH00H0H000HH0H000000000000000000
```

```
__m128i c, t, s;
c = _mm_set1_epi32(0x7f532c7f);
res[0] = _mm_and_si128(m[0], c);
c = _mm_set1_epi32(0x20400000);
t = _mm_and_si128(m[1], c);
s = _mm_srli_epi32(t, 15);
res[0] = _mm_or_si128(res[0], s);
c = _mm_set1_epi32(0x00320000);
t = _mm_and_si128(m[1], c);
s = _mm_srli_epi32(t, 2);
res[0] = _mm_or_si128(res[0], s);
c = _mm_set1_epi32(0x04050000);
t = _mm_and_si128(m[1], c);
s = _mm_slli_epi32(t, 5);
res[0] = _mm_or_si128(res[0], s);
c = _mm_set1_epi32(0x00001300);
t = _mm_and_si128(m[2], c);
res[0] = _mm_or_si128(res[0], t);

c = _mm_set1_epi32(0x0000532c);
res[1] = _mm_and_si128(m[1], c);
c = _mm_set1_epi32(0x08080000);
t = _mm_and_si128(m[1], c);
s = _mm_srli_epi32(t, 12);
res[1] = _mm_or_si128(res[1], s);
c = _mm_set1_epi32(0x00002c53);
t = _mm_and_si128(m[2], c);
res[1] = _mm_or_si128(res[1], t);
```

- 1001011000110101000010010110001101010000100101100011010100001001011000110101000010010110001101 0100001001011000110101

  **Input seed:**

  ```
  A00A0AA0 00AA0A0A 0000A00A 0AA000AA
  0B0B0000 B00B0BB0 00HH0G0H 0000H00G
  0DI000ID 0I0I0000 C00C0DC0 00DC0C0C
  0000F00F 0EF000FF 0E0E0000 00000000
  ```

  **Output seed:**

  ```
  ACDACAAC DEAACACA CEFEAFDA FAADFFAA
  IBGBIHHI BIHBGBBH 00000000 00000000
  ```

  ```
  Row  1:                    A00A0AA000AA0A0A0000A00A0AA000AA

  Ext  C:     0000000000000000C00C00C0000C0C0C
  Ext  D:                     0D00000D0000000000000D0000D00000
  Ext  E:                    000000000E0000000E0E000000000000
  ```

```
Ext  F:                                  0000F00F00F000FF0000000000000000

Row  2:                    0B0B0000B00B0BB00000000000000000

Ext  G: 000000000000000000000G000000000G
Ext  H:       000000000000000000HH000H0000H000
Ext  I:                  00I000I00I0I00000000000000000000

__m128i c, t, s;
c = _mm_set1_epi32(0xc690ac69);
res[0] = _mm_and_si128(m[0], c);
c = _mm_set1_epi32(0xa8490000);
t = _mm_and_si128(m[2], c);
s = _mm_srli_epi32(t, 15);
res[0] = _mm_or_si128(res[0], s);
c = _mm_set1_epi32(0x04200082);
t = _mm_and_si128(m[2], c);
s = _mm_slli_epi32(t, 1);
res[0] = _mm_or_si128(res[0], s);
c = _mm_set1_epi32(0x000a0200);
t = _mm_and_si128(m[3], c);
res[0] = _mm_or_si128(res[0], t);
c = _mm_set1_epi32(0x0000c490);
t = _mm_and_si128(m[3], c);
s = _mm_slli_epi32(t, 14);
res[0] = _mm_or_si128(res[0], s);

c = _mm_set1_epi32(0x0000690a);
res[1] = _mm_and_si128(m[1], c);
c = _mm_set1_epi32(0x80200000);
t = _mm_and_si128(m[1], c);
s = _mm_srli_epi32(t, 19);
res[1] = _mm_or_si128(res[1], s);
c = _mm_set1_epi32(0x108c0000);
t = _mm_and_si128(m[1], c);
s = _mm_srli_epi32(t, 13);
res[1] = _mm_or_si128(res[1], s);
c = _mm_set1_epi32(0x00000a44);
t = _mm_and_si128(m[2], c);
s = _mm_srli_epi32(t, 2);
res[1] = _mm_or_si128(res[1], s);
```

#### S7..4 Weight 56

- 111111101111111011111110111111101111111011111110111111101111111

  **Input seed:**

```
AAAAAAA0 AAAAAAA0 AAAAAAA0 AAAAAAA0
BBBBBBB0 BBBBBBB0 BBBBBBB0 IHGFEDC0
```

  **Output seed:**

```
AAAAAAAC AAAAAAAD AAAAAAAE AAAAAAAF
BBBBBBBG BBBBBBBH BBBBBBBI 00000000

Row  1:                        AAAAAAA0AAAAAAA0AAAAAAA0AAAAAAA0

Ext  C: 000000000000000000000000000000C0
Ext  D:          00000000000000000000000000000D00
Ext  E:                   0000000000000000000000000000E000
Ext  F:                            000000000000000000000000000F0000

Row  2:                        BBBBBBB0BBBBBBB0BBBBBBB000000000

Ext  G:     00000000000000000000000000G00000
Ext  H:              0000000000000000000000000H000000
Ext  I:                       000000000000000000000000I0000000
```

```
__m128i c, t, s;
c = _mm_set1_epi32(0x7f7f7f7f);
res[0] = _mm_and_si128(m[0], c);
c = _mm_set1_epi32(0x40000000);
t = _mm_and_si128(m[1], c);
s = _mm_srli_epi32(t, 23);
res[0] = _mm_or_si128(res[0], s);
c = _mm_set1_epi32(0x20000000);
t = _mm_and_si128(m[1], c);
s = _mm_srli_epi32(t, 14);
res[0] = _mm_or_si128(res[0], s);
c = _mm_set1_epi32(0x10000000);
t = _mm_and_si128(m[1], c);
s = _mm_srli_epi32(t, 5);
res[0] = _mm_or_si128(res[0], s);
c = _mm_set1_epi32(0x08000000);
t = _mm_and_si128(m[1], c);
s = _mm_slli_epi32(t, 4);
res[0] = _mm_or_si128(res[0], s);

c = _mm_set1_epi32(0x007f7f7f);
res[1] = _mm_and_si128(m[1], c);
c = _mm_set1_epi32(0x04000000);
t = _mm_and_si128(m[1], c);
s = _mm_srli_epi32(t, 19);
res[1] = _mm_or_si128(res[1], s);
c = _mm_set1_epi32(0x02000000);
t = _mm_and_si128(m[1], c);
s = _mm_srli_epi32(t, 10);
res[1] = _mm_or_si128(res[1], s);
c = _mm_set1_epi32(0x01000000);
t = _mm_and_si128(m[1], c);
s = _mm_srli_epi32(t, 1);
res[1] = _mm_or_si128(res[1], s);
```

- 1111111110110111000111111111011011100011111111101101110001111111110110111

  **Input seed:**

```
AAAAAAAA A0AA0AAA 000AAAAA AAAA0AA0
BBB000BB BBBBBBB0 BB0BBB00 0CFFFCCC
GE0GE0DG G0000000 00000000 00000000
```

  **Output seed:**

```
AAAAAAAA ADAACAAA CCCAAAAA AAAAEAAE
BBBFFFBB BBBBBBBG BBGBBBGG 00000000

Row  1:                        AAAAAAAAA0AA0AAA000AAAAAAAAA0AA0

Ext  C:           00000000000000000000000000C000CCC
Ext  D:                           000000D00000000000000000000000000
Ext  E:                                                   0E00E0000000000000000000000000000

Row  2:                        BBB000BBBBBBBBB0BB0BBB0000000000

Ext  F: 000000000000000000000000000FFF000
Ext  G:                                       G00G000GG0000000000000000000000

__m128i c, t, s;
c = _mm_set1_epi32(0x6ff8edff);
res[0] = _mm_and_si128(m[0], c);
c = _mm_set1_epi32(0xe2000000);
t = _mm_and_si128(m[1], c);
s = _mm_srli_epi32(t, 13);
res[0] = _mm_or_si128(res[0], s);
c = _mm_set1_epi32(0x00000040);
t = _mm_and_si128(m[2], c);
s = _mm_slli_epi32(t, 3);
res[0] = _mm_or_si128(res[0], s);
c = _mm_set1_epi32(0x00000012);
t = _mm_and_si128(m[2], c);
s = _mm_slli_epi32(t, 27);
res[0] = _mm_or_si128(res[0], s);

c = _mm_set1_epi32(0x003b7fc7);
res[1] = _mm_and_si128(m[1], c);
c = _mm_set1_epi32(0x1c000000);
t = _mm_and_si128(m[1], c);
s = _mm_srli_epi32(t, 23);
res[1] = _mm_or_si128(res[1], s);
c = _mm_set1_epi32(0x00000189);
t = _mm_and_si128(m[2], c);
s = _mm_slli_epi32(t, 15);
res[1] = _mm_or_si128(res[1], s);
```

- 111011111010011101001000111011111010011101001000111011111010011101001000111011111010011101001

  **Input seed:**

```
AAA0AAAA A0A00AAA 0A00A000 AAA0AAAA
B0B00BBB 0B00B000 BBB0BBBB C0C00CCC
0E00G000 GEH0HFGF H0D00DFF 0E00D000
```

  **Output seed:**

```
AAAEAAAA ADAEDAAA CACDACCC AAAEAAAA
BFBFGBBB GBFFBHGH BBBHBBBB 00000000

Row  1:             AAA0AAAAA0A00AAA0A00A000AAA0AAAA

Ext  C:     00000000000000000000000000C0C00CCC
Ext  D:    000000000000000000D00D000000D000
Ext  E:               0E0000000E000000000000000E000000

Row  2:             B0B00BBB0B00B000BBB0BBBB00000000

Ext  F: 0000000000000F0F000000FF00000000
Ext  G:             0000G000G00000G00000000000000000
Ext  H:                0000000000H0H000H000000000000000
```

```
__m128i c, t, s;
c = _mm_set1_epi32(0xf712e5f7);
res[0] = _mm_and_si128(m[0], c);
c = _mm_set1_epi32(0xe5000000);
t = _mm_and_si128(m[1], c);
s = _mm_srli_epi32(t, 8);
res[0] = _mm_or_si128(res[0], s);
c = _mm_set1_epi32(0x10240000);
t = _mm_and_si128(m[2], c);
s = _mm_srli_epi32(t, 9);
res[0] = _mm_or_si128(res[0], s);
c = _mm_set1_epi32(0x02000202);
t = _mm_and_si128(m[2], c);
s = _mm_slli_epi32(t, 2);
res[0] = _mm_or_si128(res[0], s);

c = _mm_set1_epi32(0x00f712e5);
res[1] = _mm_and_si128(m[1], c);
c = _mm_set1_epi32(0x00c0a000);
t = _mm_and_si128(m[2], c);
s = _mm_srli_epi32(t, 12);
res[1] = _mm_or_si128(res[1], s);
c = _mm_set1_epi32(0x00004110);
t = _mm_and_si128(m[2], c);
res[1] = _mm_or_si128(res[1], t);
c = _mm_set1_epi32(0x00011400);
t = _mm_and_si128(m[2], c);
s = _mm_slli_epi32(t, 3);
res[1] = _mm_or_si128(res[1], s);
```

- 10001100101001110000100011001010011100001000110010100111000010001100101001110000100011001010011100001000110010100111000010001100101001110000100011001010011100001000110010100111

  **Input seed:**

  ```
  A000AA00 A0A00AAA 0000A000 AA00A0A0
  0BBB0000 B000BB00 B0B00BBB 0000G000
  DD00D0H0 0DCH0000 D000HC00 D0H00DCC
  0000I000 EF00I0I0 0EEF0000 E000EF00
  J0J00JJJ 00000000 00000000 00000000
  ```

  **Output seed:**

  ```
  ACDDAADE AFADCAAA EEDFACCE AADEAFAD
  HBBBJHJI BJJJBBHI BIBGHBBB 00000000
  ```

```
Row  1:          A000AA00A0A00AAA0000A000AA00A0A0

Ext  C: 0000000000C0000000000C000000000CC
Ext  D:            DD00D0000D000000D0000000D0000D00
Ext  E:         00000000E00000000EE00000E000E000
Ext  F:          000000000F000000000F000000000F00

Row  2:          0BBB0000B000BB00B0B00BBB00000000

Ext  G: 00000000000000000000000000000G000
Ext  H:    000000H0000H000000000H00000H00000
Ext  I:             0000I0000000I0I000000000000000000
Ext  J:              J0J00JJJ0000000000000000000000000

__m128i c, t, s;
c = _mm_set1_epi32(0x5310e531);
res[0] = _mm_and_si128(m[0], c);
c = _mm_set1_epi32(0xc0200400);
t = _mm_and_si128(m[2], c);
s = _mm_srli_epi32(t, 9);
res[0] = _mm_or_si128(res[0], s);
c = _mm_set1_epi32(0x21010213);
t = _mm_and_si128(m[2], c);
s = _mm_slli_epi32(t, 2);
res[0] = _mm_or_si128(res[0], s);
c = _mm_set1_epi32(0x11060100);
t = _mm_and_si128(m[3], c);
s = _mm_srli_epi32(t, 1);
res[0] = _mm_or_si128(res[0], s);
c = _mm_set1_epi32(0x20080200);
t = _mm_and_si128(m[3], c);
res[0] = _mm_or_si128(res[0], t);

c = _mm_set1_epi32(0x00e5310e);
res[1] = _mm_and_si128(m[1], c);
c = _mm_set1_epi32(0x10000000);
t = _mm_and_si128(m[1], c);
s = _mm_srli_epi32(t, 9);
res[1] = _mm_or_si128(res[1], s);
c = _mm_set1_epi32(0x04100840);
t = _mm_and_si128(m[2], c);
s = _mm_srli_epi32(t, 6);
res[1] = _mm_or_si128(res[1], s);
c = _mm_set1_epi32(0x00005010);
t = _mm_and_si128(m[3], c);
s = _mm_slli_epi32(t, 3);
res[1] = _mm_or_si128(res[1], s);
c = _mm_set1_epi32(0x000000e5);
t = _mm_and_si128(m[4], c);
s = _mm_slli_epi32(t, 4);
res[1] = _mm_or_si128(res[1], s);
```

- 1011100101111101110001001011100101111101110001001011100101111101110001001011100101111101110001

  **Input seed:**

  ```
  A0AAA00A 0AAAAA0A AA000A00 A0AAA00A
  0BBBBB0B BB000B00 B0BBB00B 0FFGGF0F
  EC000C00 C0DHE00D 0HDECH0C EH000C00
  ```

  **Output seed:**

```
ACAAACEA CAAAAADA AAEDCADC AEAAACEA
HBBBBBHB BBHFFBHF BFBBBGGB 00000000

Row  1:                A0AAA00A0AAAAA0AAA000A00A0AAA00A

Ext  C:                0C000C00C00000000000C00C00000C00
Ext  D:                    0000000000D0000D00D0000000000000
Ext  E:                      E00000000000E000000E0000E0000000

Row  2:                0BBBBB0BBB000B00B0BBB00B00000000

Ext  F:  0000000000000000000000000FF00F0F
Ext  G:          000000000000000000000000000GG000
Ext  H:     00000000000H00000H000H000H000000

__m128i c, t, s;
c = _mm_set1_epi32(0x9d23be9d);
res[0] = _mm_and_si128(m[0], c);
c = _mm_set1_epi32(0x20900122);
t = _mm_and_si128(m[2], c);
res[0] = _mm_or_si128(res[0], t);
c = _mm_set1_epi32(0x00048400);
t = _mm_and_si128(m[2], c);
s = _mm_slli_epi32(t, 4);
res[0] = _mm_or_si128(res[0], s);
c = _mm_set1_epi32(0x01081001);
t = _mm_and_si128(m[2], c);
s = _mm_slli_epi32(t, 6);
res[0] = _mm_or_si128(res[0], s);

c = _mm_set1_epi32(0x009d23be);
res[1] = _mm_and_si128(m[1], c);
c = _mm_set1_epi32(0xa6000000);
t = _mm_and_si128(m[1], c);
s = _mm_srli_epi32(t, 14);
res[1] = _mm_or_si128(res[1], s);
c = _mm_set1_epi32(0x18000000);
t = _mm_and_si128(m[1], c);
s = _mm_srli_epi32(t, 6);
res[1] = _mm_or_si128(res[1], s);
c = _mm_set1_epi32(0x02220800);
t = _mm_and_si128(m[2], c);
s = _mm_srli_epi32(t, 11);
res[1] = _mm_or_si128(res[1], s);
```

#### S7..5 Weight 64

- 11111111011111111011111111011111111011111111011111111011111111011111111

  **Input seed:**

```
AAAAAAAA 0AAAAAAA A0AAAAAA AA0AAAAA
BBB0BBBB BBBB0BBB BBBBB0BB BBBBBB0B
IHGFEDC0 00000000 00000000 00000000
```

  **Output seed:**

```
AAAAAAAA CAAAAAAA ADAAAAAA AAEAAAAA
BBBFBBBB BBBBGBBB BBBBBHBB BBBBBBIB

Row  1: AAAAAAAA0AAAAAAAA0AAAAAAAA0AAAAA

Ext  C:   000000C0000000000000000000000000
Ext  D:             00000D00000000000000000000000000
Ext  E:                       0000E000000000000000000000000000

Row  2: BBB0BBBBBBBB0BBBBBBBB0BBBBBBBB0B

Ext  F: 000F0000000000000000000000000000
Ext  G:           00G00000000000000000000000000000
Ext  H:                     0H000000000000000000000000000000
Ext  I:                               I0000000000000000000000000000000

__m128i c, t, s;
c = _mm_set1_epi32(0xfbfdfeff);
res[0] = _mm_and_si128(m[0], c);
c = _mm_set1_epi32(0x00000040);
t = _mm_and_si128(m[2], c);
s = _mm_slli_epi32(t, 2);
res[0] = _mm_or_si128(res[0], s);
c = _mm_set1_epi32(0x00000020);
t = _mm_and_si128(m[2], c);
s = _mm_slli_epi32(t, 12);
res[0] = _mm_or_si128(res[0], s);
c = _mm_set1_epi32(0x00000010);
t = _mm_and_si128(m[2], c);
s = _mm_slli_epi32(t, 22);
res[0] = _mm_or_si128(res[0], s);

c = _mm_set1_epi32(0xbfdfeff7);
res[1] = _mm_and_si128(m[1], c);
c = _mm_set1_epi32(0x00000008);
t = _mm_and_si128(m[2], c);
res[1] = _mm_or_si128(res[1], t);
c = _mm_set1_epi32(0x00000004);
t = _mm_and_si128(m[2], c);
s = _mm_slli_epi32(t, 10);
res[1] = _mm_or_si128(res[1], s);
c = _mm_set1_epi32(0x00000002);
t = _mm_and_si128(m[2], c);
s = _mm_slli_epi32(t, 20);
res[1] = _mm_or_si128(res[1], s);
c = _mm_set1_epi32(0x00000001);
t = _mm_and_si128(m[2], c);
s = _mm_slli_epi32(t, 30);
res[1] = _mm_or_si128(res[1], s);
```

- 1111010111111111011001111010111111111011001111010111111111011001111010111111111011

  **Input seed:**

```
AAAA0A0A AAAAAAAA 0AA00AAA A0A0AAAA
BBBBB0BB 00BBBB0B 0BBBBBBB BB0BB00B
FDD0C0CD EDFEEFF0 CE000000 00000000
```

  **Output seed:**

```
AAAACACA AAAAAAAA CAADDAAA ADADAAAA
BBBBBEBB EEBBBBEB FBBBBBBB BBFBBFFB

Row  1:    AAAA0A0AAAAAAAAA0AA00AAAA0A0AAAA

Ext  C:    0000C0C000000000C000000000000000
Ext  D:                      0DD0000D0D0000000000000000000000

Row  2:    BBBBB0BB00BBBB0B0BBBBBBBBB0BB00B

Ext  E: 00000000E00EE0000E000000000000000
Ext  F:                    F000000000F00FF00000000000000000

__m128i c, t, s;
c = _mm_set1_epi32(0xf5e6ffaf);
res[0] = _mm_and_si128(m[0], c);
c = _mm_set1_epi32(0x00010050);
t = _mm_and_si128(m[2], c);
res[0] = _mm_or_si128(res[0], t);
c = _mm_set1_epi32(0x00000286);
t = _mm_and_si128(m[2], c);
s = _mm_slli_epi32(t, 18);
res[0] = _mm_or_si128(res[0], s);

c = _mm_set1_epi32(0x9bfebcdf);
res[1] = _mm_and_si128(m[1], c);
c = _mm_set1_epi32(0x00021900);
t = _mm_and_si128(m[2], c);
s = _mm_srli_epi32(t, 3);
res[1] = _mm_or_si128(res[1], s);
c = _mm_set1_epi32(0x00006401);
t = _mm_and_si128(m[2], c);
s = _mm_slli_epi32(t, 16);
res[1] = _mm_or_si128(res[1], s);
```

- 1110111110100111010010001110111110100111010010001110111110100111010010001110111110100111010010 00111011111

  **Input seed:**

```
AAA0AAAA A0A00AAA 0A00A000 AAA0AAAA
B0B00BBB 0B00B000 BBB0BBBB B0B00BBB
0G00H000 HDE0DGGC H0D00EDC 0G00G000
IIF0FIFI I0000000 00000000 00000000
```

  **Output seed:**

```
AAACAAAA ADACDAAA EADFAFDF AAAEAAAA
BGBIIBBB IBIIBGGH BBBHBBBB BGBHGBBB
```

```
Row  1:             AAA0AAAAA0A00AAA0A00A000AAA0AAAA

Ext  C: 000000000000000C0000000C00000000
Ext  D:             000000000D00D00000D000D000000000
Ext  E:                   0000000000E0000000000E0000000000
Ext  F:                              00F0F0F0000000000000000000000000

Row  2:             B0B00BBB0B00B000BBB0BBBBB0B00BBB

Ext  G:             0G00000000000GG0000000000G00G000
Ext  H:                        0000H000H0000000H000000000000000
Ext  I:                II000I0II00000000000000000000000
```

```
__m128i c, t, s;
c = _mm_set1_epi32(0xf712e5f7);
res[0] = _mm_and_si128(m[0], c);
c = _mm_set1_epi32(0x00808000);
t = _mm_and_si128(m[2], c);
s = _mm_srli_epi32(t, 12);
res[0] = _mm_or_si128(res[0], s);
c = _mm_set1_epi32(0x00441200);
t = _mm_and_si128(m[2], c);
res[0] = _mm_or_si128(res[0], t);
c = _mm_set1_epi32(0x00200400);
t = _mm_and_si128(m[2], c);
s = _mm_slli_epi32(t, 6);
res[0] = _mm_or_si128(res[0], s);
c = _mm_set1_epi32(0x00000054);
t = _mm_and_si128(m[3], c);
s = _mm_slli_epi32(t, 17);
res[0] = _mm_or_si128(res[0], s);

c = _mm_set1_epi32(0xe5f712e5);
res[1] = _mm_and_si128(m[1], c);
c = _mm_set1_epi32(0x12006002);
t = _mm_and_si128(m[2], c);
res[1] = _mm_or_si128(res[1], t);
c = _mm_set1_epi32(0x00010110);
t = _mm_and_si128(m[2], c);
s = _mm_slli_epi32(t, 11);
res[1] = _mm_or_si128(res[1], s);
c = _mm_set1_epi32(0x000001a3);
t = _mm_and_si128(m[3], c);
s = _mm_slli_epi32(t, 3);
res[1] = _mm_or_si128(res[1], s);
```

- 110101111011000100000010000011010111101100010000001000001101011110110001000000100000110101111011
  00010000001000001101011110110001000000100000110101111011

  **Input seed:**

  ```
  AA0A0AAA A0AA000A 00000A00 000AA0A0
  BBBB0BB0 00B00000 B00000BB 0B0BBBB0
  GC000G00 000C0000 0GC0G0GC CC0GC000
  F00000E0 0000FF0E 0EFHE0DD 000F0000
  0J00000I I0J0IIII 0II00000 00000000
  ```

  **Output seed:**

  ```
  AACAFAAA AEAACDDA FFECEAFE CCCAACAF
  BBBBGBBI IGBJIIII BIIHJGBB GBGBBBBG
  ```

```
Row  1:          AA0A0AAAA0AA000A00000A00000AA0A0

Ext  C:           0C000000000C000000C0000CCC00C000
Ext  D: 0000000000000000000000DD00000000
Ext  E:             000000E00000000E0E00E00000000000
Ext  F:              F00000000000FF0000F00000000F0000

Row  2:          BBBB0BB000B00000B00000BB0B0BBBB0

Ext  G:              G0000G00000000000G00G0G0000G0000
Ext  H:          0000000000000000000H000000000000
Ext  I:          0000000II000IIII0II0000000000000
Ext  J:                    0J00000000J0000000000000000000000

__m128i c, t, s;
c = _mm_set1_epi32(0x58208deb);
res[0] = _mm_and_si128(m[0], c);
c = _mm_set1_epi32(0x13840802);
t = _mm_and_si128(m[2], c);
s = _mm_slli_epi32(t, 1);
res[0] = _mm_or_si128(res[0], s);
c = _mm_set1_epi32(0x00c00000);
t = _mm_and_si128(m[3], c);
s = _mm_srli_epi32(t, 9);
res[0] = _mm_or_si128(res[0], s);
c = _mm_set1_epi32(0x00128040);
t = _mm_and_si128(m[3], c);
s = _mm_slli_epi32(t, 3);
res[0] = _mm_or_si128(res[0], s);
c = _mm_set1_epi32(0x08043001);
t = _mm_and_si128(m[3], c);
s = _mm_slli_epi32(t, 4);
res[0] = _mm_or_si128(res[0], s);

c = _mm_set1_epi32(0x7ac1046f);
res[1] = _mm_and_si128(m[1], c);
c = _mm_set1_epi32(0x08520021);
t = _mm_and_si128(m[2], c);
s = _mm_slli_epi32(t, 4);
res[1] = _mm_or_si128(res[1], s);
c = _mm_set1_epi32(0x00080000);
t = _mm_and_si128(m[3], c);
res[1] = _mm_or_si128(res[1], t);
c = _mm_set1_epi32(0x0006f180);
t = _mm_and_si128(m[4], c);
res[1] = _mm_or_si128(res[1], t);
c = _mm_set1_epi32(0x00000402);
t = _mm_and_si128(m[4], c);
s = _mm_slli_epi32(t, 10);
res[1] = _mm_or_si128(res[1], s);
```
